# Ena/VASP processive elongation is modulated by avidity on actin filaments bundled by the filopodia crosslinker fascin

**DOI:** 10.1101/386961

**Authors:** Alyssa J. Harker, Harshwardhan H. Katkar, Tamara C. Bidone, Fikret Aydin, Gregory A. Voth, Derek A. Applewhite, David R. Kovar

**Author notes:** Contributed equally to this work (Harshwardhan H. Katkar, Tamara C. Bidone). Address correspondence to: David R. Kovar The University of Chicago 920 East 58^th^ Street CLSC Suite 212 Chicago, IL 60637.

## Abstract

Ena/VASP are tetrameric assembly factors that bind F-actin barbed ends continuously while increasing their elongation rate within dynamic bundled networks such as filopodia. We used single-molecule TIRFM and developed a kinetic model to dissect Ena/VASP’s processive mechanism on bundled filaments. Notably, Ena/VASP’s processive run length increases with the number of both bundled filaments and Ena arms, revealing avidity facilitates enhanced processivity. Moreover, Ena tetramers form more filopodia than mutant dimer and trimers in *Drosophila* culture cells. Finally, enhanced processivity on trailing barbed ends of bundled filaments is an evolutionarily conserved property of Ena/VASP homologs and is specific to fascin-bundled filaments. These results demonstrate that Ena tetramers are tailored for enhanced processivity on fascin bundles and avidity of multiple arms associating with multiple filaments is critical for this process. Furthermore, we discovered a novel regulatory mechanism whereby bundle size and bundling protein specificity control activities of a processive assembly factor.

## INTRODUCTION

Many important cellular functions depend on formation of actin cytoskeleton networks at the correct time and location with specific architectures and dynamics (Campellone and Welch, 2010; Pollard and Cooper, 2009). For example, filopodia are filamentous actin (F-actin)-rich finger-like protrusions that elongate from the lamellipodium, a dense, branched F-actin network kept short by capping protein (Pollard and Borisy, 2003) at the cell periphery. Filopodia are important for cell motility and environment sensing. Filopodial actin filaments are assembled by actin elongation factors such as formins and Enabled/vasodilator-stimulated phosphoprotein (Ena/VASP) (Mattila and Lappalainen, 2008). During filopodia initiation Ena/VASP localizes to the edge of the lamellipodium where it competes with capping protein for barbed ends (Applewhite et al., 2007; Barzik et al., 2005; Bear et al., 2002; Bear and Gertler, 2009; Svitkina et al., 2003; Winkelman et al., 2014), and then facilitates generation of long, straight filaments by remaining processively associated with barbed ends and increasing their elongation rate 2- to 7-fold (Breitsprecher et al., 2011, 2008; Brühmann et al., 2017; Hansen and Mullins, 2010; Pasic et al., 2008; Winkelman et al., 2014). The 10-30 filaments in filopodia are bundled primarily by fascin, a globular crosslinking protein containing β-trefoil domains (Jansen et al., 2011; Vignjevic et al., 2006). Fascin bundles are composed of parallel filaments with narrow spacing, between 8-10 nm (Cant et al., 1994; Edwards and Bryan, 1995; Jansen et al., 2011; Yang et al., 2013). Ena/VASP continues to localize to the tips of mature filopodia, where fascin-bundled filaments ultimately are the same length (Faix and Rottner, 2006; Gupton and Gertler, 2007), presumably assuring uniform thickness of filopodia required for protrusive force (Svitkina et al., 2003; Winkelman et al., 2014).

Ena/VASP is a multidomain homotetramer with homologs in all metazoan cells (Sebé-Pedrós et al., 2013). A few Ena/VASP homologs have been biochemically characterized including human VASP (Bachmann et al., 1999; Breitsprecher et al., 2008; Chereau and Dominguez, 2006; Hansen and Mullins, 2010; Pasic et al., 2008), *Drosophila* Enabled (Winkelman et al., 2014), and *Dictyostelium* VASP (Breitsprecher et al., 2008). Ena/VASP proteins contain two conserved Ena/VASP homology domains, EVH1 and EVH2 (Figure 1A). The N-terminus EVH1 domain is important for cellular localization and binds to proteins with FPPPP (FP4) repeats, such as lamellipodin, zyxin, and formin (Ball et al., 2001; Bilancia et al., 2014). The C-terminus EVH2 domain consists of three smaller subdomains: G-actin binding domain (GAB) (Bachmann et al., 1999; Ferron et al., 2007), F-actin binding domain (FAB) (Dominguez and Holmes, 2011), and a C-terminal coiled-coil tetramerization domain (Bachmann et al., 1999; Kuhnel et al., 2004). Between the EVH1 and EVH2 domains there is a poly-proline rich region that binds profilin as well as SH3 domains (Ferron et al., 2007; Hansen and Mullins, 2010).

**Figure 1:**
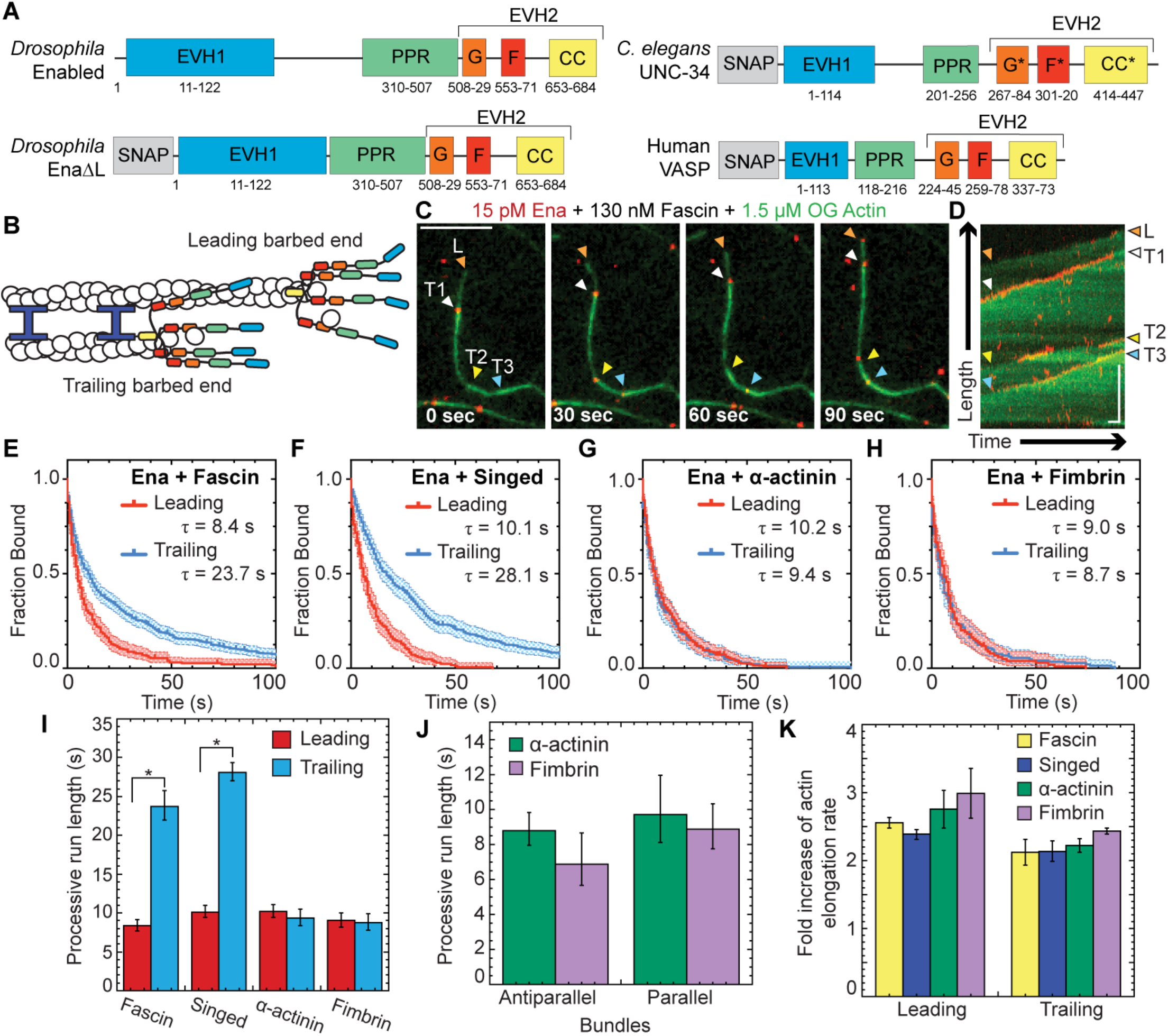
Ena has enhanced processivity on F-actin bundles formed specifically by fascin. (A) Ena/VASP domain organization and constructs used for Ena, UNC-34, and VASP: Self-labeling tag (SNAP), Ena/VASP homology domain 1 (EVH1), polyproline region (PPR), Ena/VASP homology domain 2 (EVH2) includes G-actin binding domain (G), F-actin binding domain (F), coiled coil region (CC). *Putative domain. Two-color TIRFM visualization of 1.5 μM Mg-ATP-actin (15% Oregon green-actin) with 15 pM SNAP(549)-EnaΔL and unlabeled 130 nM human fascin, 250 nM fly fascin Singed, 125 nM α-actinin, or 100 nM fimbrin. (B) Cartoon Ena/VASPs bound to leading and trailing barbed ends in a fascin bundle. (C and D) Representative experiment of OG-actin with SNAP(549)-EnaΔL and fascin. Arrows indicate leading (orange), 1^st^ trailing (white), 2^nd^ trailing (yellow) and 3^rd^ trailing (blue) barbed ends. (C) Merged time-lapse micrographs. Scale bar, 5 μm. (D) Merged kymograph of filament length (scale bar, 5 μm) over time (time bar, 10 s). (E-H) Kaplan-Meier curves representing average processive run lengths (τ) for Ena with (E) fascin, (F) Singed, (G) α-actinin, or (H) fimbrin on leading (red) and trailing (blue) barbed ends. Error bars, 95% CI. n ≥ 127. (I) Average processive run lengths for leading (red) and trailing (blue) barbed ends shown in E-H for 2-filament bundles with fascin, Singed, α-actinin, or fimbrin. P values (*<0.0001). Error bars, 95% CI. (J) Average processive run lengths for antiparallel and parallel 2-filament α-actinin (green) or fimbrin (purple) bundles. Error bars, 95% CI. n ≥ 64. (K) Fold increase of barbed end elongation rates of Ena on fascin (yellow), Singed (blue), α-actinin (green), or fimbrin (purple) bundled filaments. Error bars, SEM. n ≥ 5 barbed ends from at least 2 movies.

In addition to the leading edge and tips of filopodia, Ena/VASP proteins also localize to focal adhesions and stress fibers (Brindle et al., 1996; Reinhard et al., 1992), which are composed of filaments crosslinked by CH domain superfamily crosslinkers, fimbrin/plastin and α-actinin. Fimbrin also localizes to the lamellipodia and base of filopodia, and it bundles both parallel and antiparallel filaments with narrow spacing (10-12 nm), similar to fascin (Hanein et al., 1998). In comparison, α-actinin bundles filaments of mixed polarity with much wider spacing (30-36 nm) (Sjöblom et al., 2008).

We previously discovered that for F-actin bundles made by human fascin, *Drosophila* Enabled (Ena) remains processively associated with trailing barbed ends (shorter filaments) ~3-fold longer than leading barbed ends (longest filament) (Figure 1B) (Winkelman et al., 2014). We hypothesized that Ena’s increased processivity on trailing barbed ends contributes to robust filopodia formation through promoting growth of these shorter filaments by prolonged protection against capping protein and increased elongation rate. Trailing barbed ends are thereby allowed to catch up to leading barbed ends ensuring mature filopodia with bundled filaments of uniform length. However, the underlying molecular mechanisms that facilitate Ena’s enhanced processivity on bundled filaments remain unclear.

We used a combination of in vitro reconstitution with single-molecule multi-color total internal reflection fluorescence microscopy (TIRFM), kinetic modeling, and analysis of *Drosophila* culture cells to characterize the dynamics and function of processive elongation of single and bundled filaments by multiple Ena/VASP homologs including Ena, human VASP, and *C. elegans* UNC-34. We discovered that enhanced processivity on trailing barbed ends is specific to fascin bundles and is positively correlated with the number of filaments in a bundle as well as the number of Ena monomers, or ‘arms’, available to bind nearby filaments. We also observed that Ena tetramers are more efficient at forming filopodia in *Drosophila* culture cells compared to Ena dimers and trimers. Together, our experiments and simulations inform our mechanistic understanding of Ena/VASP on single and bundled filaments and demonstrate that avidity of multiple filaments within fascin bundles and multiple Ena arms leads to increased processivity of tetrameric Ena on trailing barbed ends.

## RESULTS

### Ena is more processive on trailing barbed ends of both human and fly fascin (Singed) bundles

To understand what features are important for *Drosophila* Ena’s enhanced processivity on trailing barbed ends within human fascin bundles (Figure 1B) (Winkelman et al., 2014), we first tested if a different fascin homolog also facilitates enhanced residence times. We used two-color TIRFM to directly visualize the assembly of 1.5 μM Mg-ATP-actin monomers (15% Oregon green-labeled) with 15 pM fluorescently labeled SNAP(549)-EnaΔL (referred to as Ena) (Figure 1A) and human fascin or fly fascin, Singed. TIRFM allows direct visualization of individual Ena molecule dynamics on single and bundled actin filament barbed ends. Ena processive run lengths were measured for leading and single filament barbed ends (collectively referred to as leading) as well as trailing barbed ends (Figure 1C-D, Movie 1). Kaplan Meier survival curves were calculated from individual Ena processive runs (Figure 1E-F), revealing that Ena remains associated with trailing barbed ends (τ_fascin_ = 23.7 s, τ_Singed_ = 28.1 s) ~3-fold longer than leading barbed ends (τ_fascin_ = 8.4 s, τ_Singed_ = 10.1 s) for both human fascin and fly Singed (Figure 1I, Table 1), consistent with our previous findings (Winkelman et al., 2014). Therefore, enhancement of Ena’s processive elongation on trailing barbed ends is not specific to a particular fascin homolog.

**Table 1.** Comparison of Ena/VASP proteins’ residence time on various bundled F-actin, Related to Results and Discussion

| Ena/<br>VASP | Bundling<br>Protein | Leading<sup>a</sup> (s) | Trailing<sup>a</sup> (s) | L/T<br>p-value<sup>b</sup> | 1 fil. <sup>c</sup><br>(s) | 2 fil. <sup>c</sup><br>(s) | ≥ 3 fil. <sup>c</sup><br>(s) | 1/2/≥ 3<br>p-value<sup>d</sup> |
| --- | --- | --- | --- | --- | --- | --- | --- | --- |
| Ena<br>Tetramer | Fascin | 8.4<br>[7.7, 9.1]<br>(254) | 23.7<br>[22.0, 25.8]<br>(511) | <.0001 | 8.9<br>[7.5, 10.6]<br>(107) | 16.8<br>[14.3, 19.7]<br>(201) | 26.0<br>[23.7, 28.6]<br>(308) | <.0001 |
| Ena<br>Tetramer | Singed | 10.1<br>[9.4, 11.0]<br>(184) | 28.1<br>[27.0, 29.4]<br>(328) | <.0001 | 10.0<br>[9.2, 10.8]<br>(98) | 21.7<br>[20.4, 23.3]<br>(155) | 32.3<br>[31.0, 33.6]<br>(176) | <.0001 |
| Ena<br>Tetramer | α-actinin | 10.2<br>[9.5, 11.1]<br>(284) | 9.4<br>[8.4, 10.5]<br>(176) | 0.64 | 9.1<br>[8.3, 10.2]<br>(165) | 8.9<br>[7.7, 10.6]<br>(116) | 8.7<br>[7.8, 9.9]<br>(60) | 0.56 |
| Ena<br>Tetramer | Fimbrin | 9.0<br>[8.2, 10.0]<br>(127) | 8.7<br>[7.7, 10.0]<br>(183) | 0.91 | 8.2<br>[7.1, 9.8]<br>(63) | 7.6<br>[6.6, 9.1]<br>(121) | 10.2<br>[8.6, 12.6]<br>(64) | 0.33 |
| VASP<br>Tetramer | Fascin | 1.2<br>[0.9, 1.6]<br>(213) | 3.3<br>[3.1, 3.5]<br>(348) | <.0001 | 1.0<br>[0.7, 1.8]<br>(123) | 2.6<br>[2.4, 2.9]<br>(207) | 4.2<br>[3.9, 4.4]<br>(143) | <.0001 |
| VASP<br>Tetramer | Singed | 1.0<br>[0.7, 1.5]<br>(187) | 3.5<br>[3.2, 3.8]<br>(463) | <.0001 | 0.9<br>[0.6, 1.6]<br>(118) | 2.8<br>[2.6, 3.2]<br>(224) | 4.1<br>[3.7, 4.7]<br>(220) | <.0001 |
| UNC-34<br>Tetramer | Fascin | 1.7<br>[1.4, 2.2]<br>(82) | 3.7<br>[3.4, 4.0]<br>(266) | <.0001 | 1.2<br>[0.9, 1.8]<br>(65) | 2.9<br>[2.7, 3.3]<br>(123) | 3.9<br>[3.5, 4.5]<br>(144) | <.0001 |
| Ena Trimer | Fascin | 6.2<br>[5.6, 6.9]<br>(322) | 9.8<br>[9.2, 10.4]<br>(299) | <.0001 | 5.3<br>[4.7, 6.1]<br>(151) | 8.9<br>[8.3, 9.6]<br>(206) | 11.2<br>[10.2, 12.6]<br>(93) | <.0001 |
| Ena Dimer | Fascin | 1.3<br>[1.0, 1.7]<br>(376) | 1.8<br>[1.5, 2.4]<br>(418) | 0.01 | 1.2<br>[0.9, 1.9]<br>(197) | 1.5<br>[1.2, 2.0]<br>(261) | 2.5<br>[1.9, 3.8]<br>(122) | 0.03 |

<sup>a</sup> Values of average processive lifetime (s) [95%CI] (n) where n is the number of Ena/VASP binding events measured in at least three movies for Leading or Trailing barbed ends.
<sup>b</sup> Log Rank p-value comparing Leading and Trailing average processive lifetime.
<sup>c</sup> Values of average processive lifetime (s) [95%CI] (n) where n is the number of Ena/VASP binding events measured in at least three movies for 1 filament (fil.), 2 filaments, or greater than or equal to 3 filaments barbed ends.
<sup>d</sup> Log Rank p-value comparing 1 filament, 2 filaments, or greater than or equal to 3 filaments average processive lifetime.

### Ena’s residence time is not enhanced on trailing barbed ends of fimbrin and α-actinin bundles

To determine if diverse bundle architectures are similarly sufficient to enhance Ena’s processivity on trailing barbed ends, we tested the effect of bundling proteins with distinct properties (fimbrin and α-actinin, see introduction). First, we measured elongation rates of Ena-bound leading and trailing barbed ends of filaments bundled by human fascin, fly fascin Singed, α-actinin, or fimbrin. Two-color TIRFM visualization of control and Ena-bound barbed ends revealed a similar fold increase in Ena-mediated actin elongation for leading (~2.2- to 3-fold) and trailing (~2- to 2.5-fold) barbed ends with all four bundling proteins (Figure 1K, Tables 2-3). Therefore, Ena’s barbed end elongation enhancement is bundling protein independent.

**Table 2:**
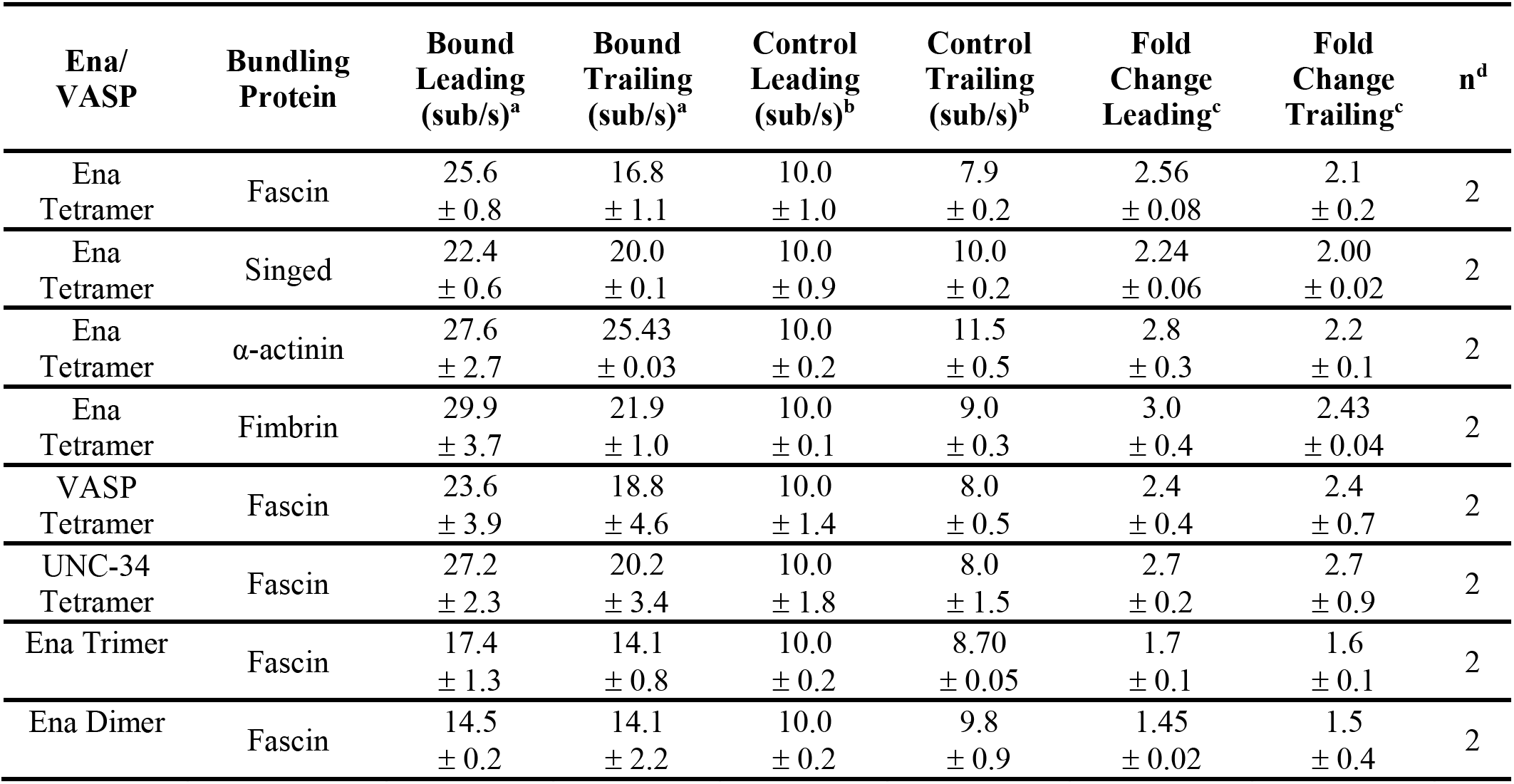
Comparison of actin elongation rates with and without (control) Ena/VASP bound, Related to Results and Discussion

| Ena/<br>VASP | Bundling<br>Protein | Bound<br>Leading<br>(sub/s)<sup>a</sup> | Bound<br>Trailing<br>(sub/s)<sup>a</sup> | Control<br>Leading<br>(sub/s)<sup>b</sup> | Control<br>Trailing<br>(sub/s)<sup>b</sup> | Fold<br>Change<br>Leading<sup>c</sup> | Fold<br>Change<br>Trailing<sup>c</sup> | n<sup>d</sup> |
| --- | --- | --- | --- | --- | --- | --- | --- | --- |
| Ena<br>Tetramer | Fascin | 25.6<br>± 0.8 | 16.8<br>± 1.1 | 10.0<br>± 1.0 | 7.9<br>± 0.2 | 2.56<br>± 0.08 | 2.1<br>± 0.2 | 2 |
| Ena<br>Tetramer | Singed | 22.4<br>± 0.6 | 20.0<br>± 0.1 | 10.0<br>± 0.9 | 10.0<br>± 0.2 | 2.24<br>± 0.06 | 2.00<br>± 0.02 | 2 |
| Ena<br>Tetramer | α-actinin | 27.6<br>± 2.7 | 25.43<br>± 0.03 | 10.0<br>± 0.2 | 11.5<br>± 0.5 | 2.8<br>± 0.3 | 2.2<br>± 0.1 | 2 |
| Ena<br>Tetramer | Fimbrin | 29.9<br>± 3.7 | 21.9<br>± 1.0 | 10.0<br>± 0.1 | 9.0<br>± 0.3 | 3.0<br>± 0.4 | 2.43<br>± 0.04 | 2 |
| VASP<br>Tetramer | Fascin | 23.6<br>± 3.9 | 18.8<br>± 4.6 | 10.0<br>± 1.4 | 8.0<br>± 0.5 | 2.4<br>± 0.4 | 2.4<br>± 0.7 | 2 |
| UNC-34<br>Tetramer | Fascin | 27.2<br>± 2.3 | 20.2<br>± 3.4 | 10.0<br>± 1.8 | 8.0<br>± 1.5 | 2.7<br>± 0.2 | 2.7<br>± 0.9 | 2 |
| Ena Trimer | Fascin | 17.4<br>± 1.3 | 14.1<br>± 0.8 | 10.0<br>± 0.2 | 8.70<br>± 0.05 | 1.7<br>± 0.1 | 1.6<br>± 0.1 | 2 |
| Ena Dimer | Fascin | 14.5<br>± 0.2 | 14.1<br>± 2.2 | 10.0<br>± 0.2 | 9.8<br>± 0.9 | 1.45<br>± 0.02 | 1.5<br>± 0.4 | 2 |

<sup>a</sup> Normalized actin elongation rate (sub/s) of Ena/VASP bound Leading or Trailing barbed ends to Control Leading.
<sup>b</sup> Normalized actin elongation rate (sub/s) of Ena/VASP free Leading or Trailing barbed ends to Control Leading.
<sup>c</sup> Fold change in actin elongation rate of Ena/VASP bound over Ena/VASP free Leading or Trailing barbed ends.
<sup>d</sup> n is the number of movies analyzed. Each movie had at least five filaments with at least 50 length measurements for each movie.

**Table 3:** p-values for comparisons of fold change in actin elongation rate with Ena on different bundling proteins for both leading and trailing filaments.

| **Leading/**<br>**Trailing<sup>a</sup>** | **Fascin** | **Singed** | **α-actinin** | **Fimbrin** |
| --- | --- | --- | --- | --- |
| **Fascin** | 1 / 1 | 0.3 / 1 | 0.6 / 0.7 | 0.4 / 0.3 |
| **Singed** | 0.3 / 1 | 1 / 1 | 0.4 / 0.7 | 0.3 / 0.3 |
| **α-actinin** | 0.6 / 0.7 | 0.4 / 0.7 | 1 / 1 | 0.7 / 0.3 |
| **Fimbrin** | 0.4 / 0.3 | 0.3 / 0.3 | 0.7 / 0.3 | 1 / 1 |

<sup>a</sup> p-values from student's two-tailed t-test with unequal variance between fold change of actin elongation rates when Ena is bound to the Leading or Trailing barbed end.

Conversely, Ena’s enhanced processivity on trailing barbed ends is specific to fascin bundles. The average processive run length on leading barbed ends with all four bundling proteins is similar, ~10 sec (Figure 1I). However, there is no enhancement of Ena’s average residence time on trailing barbed ends of α-actinin (τ = 9.4 s) or fimbrin (τ = 8.7 s) bundles (Figure 1G-I, Table 1). Therefore, F-actin bundling proteins are not universally sufficient to enhance Ena’s processivity on trailing barbed ends. Although fascin exclusively forms parallel bundles, α-actinin and fimbrin form bundles composed of filaments with mixed polarities. We therefore compared Ena’s residence time on trailing barbed ends in parallel and antiparallel two-filament bundles. For both fimbrin and α-actinin bundles, the average residence time for trailing parallel and antiparallel barbed ends is equivalent; thus, neither bundler enhances Ena’s processivity (Figure 1J, Table 4). Therefore, neither ‘fascin-like’ filament spacing (8-10 nm) nor polarity (parallel) of actin filaments within bundles is sufficient to facilitate increased processivity on trailing barbed ends. Given that Ena’s ~3-fold enhancement of processivity on trailing barbed ends is specific to fascin, different bundling proteins could regulate Ena’s specific activity for different F-actin networks.

**Table 4:** Comparison of Ena/VASP proteins’ residence time on parallel and antiparallel bundled F-actin, Related to Results and Discussion

| Ena/<br>VASP | Bundling<br>Protein | Parallel<sup>a</sup> (s) | Anti-parallel<sup>a</sup> (s) | A/P<br>p-value<sup>b</sup> |
| --- | --- | --- | --- | --- |
| Ena<br>Tetramer | Fascin | 16.8<br>[14.3, 19.7]<br>(201) | N/A | N/A |
| Ena<br>Tetramer | Singed | 21.7<br>[20.4, 23.3]<br>(155) | N/A | N/A |
| Ena<br>Tetramer | α-actinin | 9.7<br>[8.1, 12.0]<br>(77) | 8.8<br>[8.0, 9.8]<br>(90) | 0.52 |
| Ena<br>Tetramer | Fimbrin | 8.9<br>[7.7, 10.4]<br>(64) | 6.9<br>[5.7, 8.7]<br>(106) | 0.53 |

<sup>a</sup> Values of average processive lifetime (s) [95%CI] (n) where n is the number of Ena/VASP binding events measured in at least three movies for Parallel or Anti-parallel barbed ends on 2 filament bundles. Fascin results are equal to the values in 2 filaments because fascin only makes parallel bundles.
<sup>b</sup> Log Rank p-value comparing average processive lifetime trailing barbed ends in parallel and antiparallel bundles.

### Ena’s processive run length increases with bundle size

Filopodia are composed of ~10-30 actin filaments bundled by fascin (Faix and Rottner, 2006; Svitkina et al., 2003), suggesting an avidity mechanism where enhanced processivity depends on Ena simultaneously associating with a barbed end and sides of neighboring filaments. To test whether the number of filaments in a fascin bundle positively correlates with processive run length, we determined the dependence of Ena’s enhanced processivity on fascin bundle size (Figure 2A). Average run lengths on trailing barbed ends (Figure 1E-F) was thereby parsed into 2-filament bundles or 3- or more filament bundles for both human and fly fascin (Figure 2B-D, Table 1). Ena’s average residence time on trailing barbed ends of a 2-filament bundle (τ_fascin_ = 16.8 s, τ_Singed_ = 21.7 s) is ~2-fold longer than on single filament barbed ends (τ_fascin_ = 8.9 s, τ_Singed_ = 10.0 s). Furthermore, there is an additional ~1.5-fold increase in processivity when Ena is bound to trailing barbed ends of 3- or more filament bundles (τ_fascin_ = 26.0 s, τ_Singed_ = 32.2 s) (Figure 2D). Therefore, consistent with an avidity effect, Ena’s processivity increases with the number of fascin-bundled filaments.

**Figure 2:**
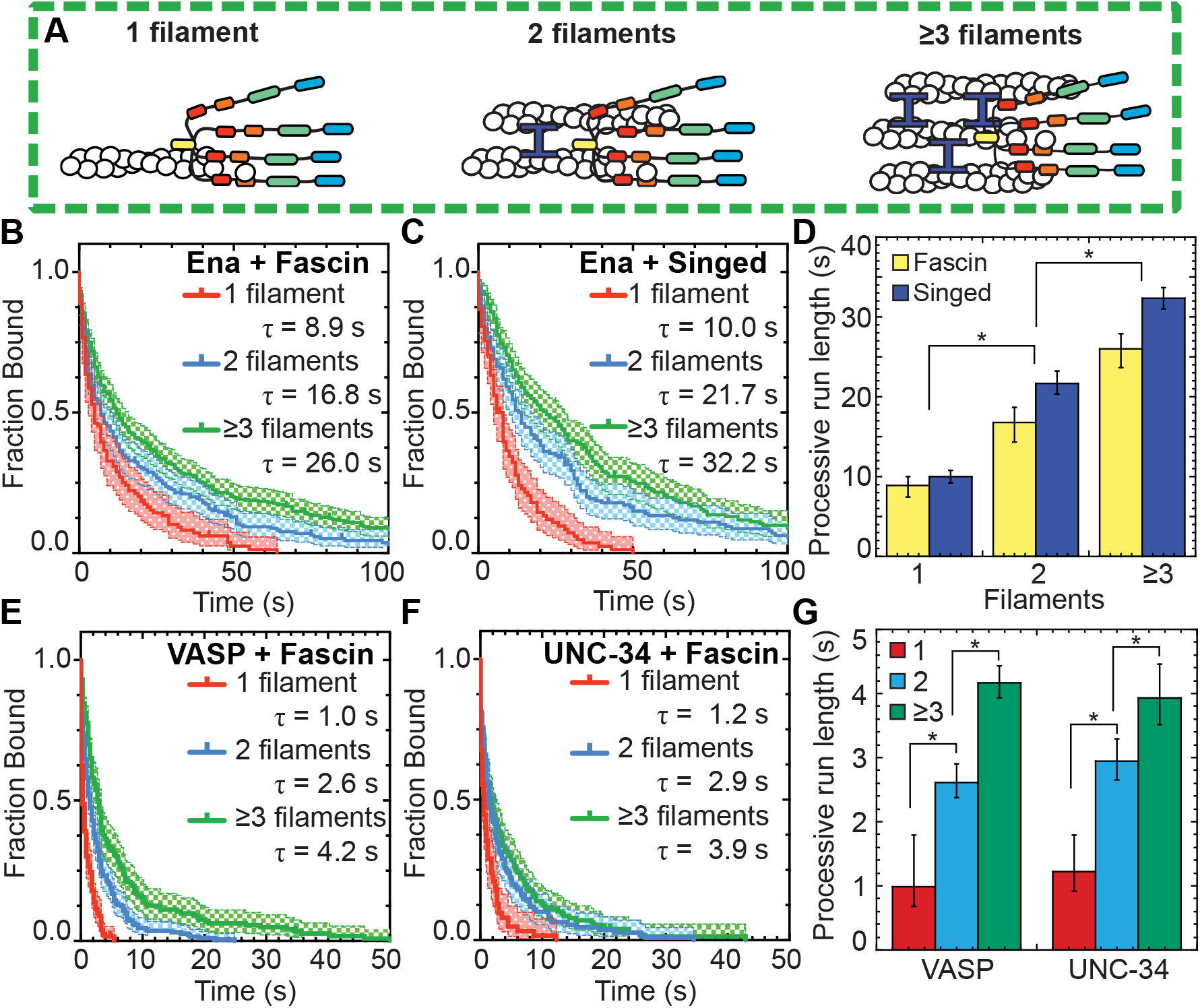
Ena/VASP’s processive run length increases with the number of filaments in a fascin bundle. (A) Cartoons of Ena/VASP on a single filament and 2- and 3-filament fascin bundles. (B-G) Two-color TIRFM visualization of 1.5 μM Mg-ATP-actin (15% Oregon green-actin) with fly SNAP(549)-EnaΔL (red), human SNAP(549)-VASP or wo)m SNAP(549)-UNC-34 and unlabeled 130 nM human fascin or 250 nM Singed as indicated. (B and C) Kaplan-Meier curves representing average processive run lengths (τ) for 15 pM Ena with (B) fascin or (C) Singed on single filaments (red), or bundles with 2 (blue) and ≥3 (green) filaments. Error bars, 95% CI. n ≥ 98. (D) Average processive run lgngthi for increasing number of filaments in fascin (yellow) or Singed (blue) bundles shown in B and C. Error bars, 95%) CI. P values (*<0.0001). (E and F) Kaplan-Meier curves representing run lengths (Δ) for (E) 25 pM VASP or (F) 1e pM UNC-34 with fascis on single filaments (red), or bundles with 2 (blue) anP ≥3 (green) filaments. Error bars, 95% CI. n ≥ 60. (G) VASP and UNC-34 average processive run lengths for increasing number of filaments in fascin bundles shown in E and F. Error bars, 95% CI. P values (*<0.0001).

### Human VASP and worm UNC-34 also have enhanced processive properties on fascin bundles

To determine whether enhanced processivity on fascin-bundled trailing filament barbed ends is conserved among Ena/VASP family members, we extended our analysis to human VASP and worm UNC-34 (Figure 1A). Human VASP is a well-characterized Ena/VASP protein (Bachmann et al., 1999; Breitsprecher et al., 2008; Chereau and Dominguez, 2006; Hansen and Mullins, 2010; Pasic et al., 2008), whereas UNC-34 had not yet been biochemically characterized in vitro despite multiple in vivo studies (Fleming et al., 2010; Havrylenko et al., 2015; Sheffield et al., 2007).

For our initial characterization of the three homologs, we measured the affinity for barbed ends and effect on actin elongation for Ena, VASP and UNC-34. Initially, the effect of Ena/VASP homologs on actin elongation rates and their apparent affinity (Kd, app) for barbed ends was determined by single-color TIRFM visualization of spontaneous assembly of 1.5 μM Mg-ATP-actin (15% Oregon Green) over a range of concentrations for each unlabeled Ena/VASP homolog (Figure 2 – figure supplement 1A-F). All three Ena/VASP homologs increase actin elongation by a similar amount, ~1.6- to ~2.7-fold, at or near saturating conditions, but have somewhat varying affinities for actin filament barbed ends ranging from 3.2 nM (Ena) to 6.7 nM (UNC-34) to 12.2 nM (VASP) (Figure 2 – figure supplement 1F). Likewise, bulk seeded pyrene actin assembly assays also show that all three Ena/VASP homologs increase actin elongation rates by similar amounts, and fits of assembly rate over a range of Ena/VASP concentrations revealed apparent affinities for barbed ends ranging from 0.7 nM (Ena) to 10.2 nM (UNC-34) to 10.8 nM (VASP) (Figure 2 – figure supplement 1G-H). We then used two-color TIRFM visualizations of red-labeled Ena, VASP, and UNC-34 on fascin bundles to measure actin elongation rates of Ena/VASP-bound leading and trailing barbed ends (Figure 2 – figure supplement 1I, Movie 2). All three Ena/VASP homologs similarly increase actin elongation ~2- to 3-fold on both leading and trailing barbed ends (Figure 2 – figure supplement 1J, Tables 3,5). Enhancement of actin elongation rates by Ena and VASP are similar to previously reported values (Brühmann et al., 2017; Hansen and Mullins, 2010; Winkelman et al., 2014) and the actin elongation properties of UNC-34 are in good agreement with the other homologs. Therefore, though Ena, VASP, and UNC-34 vary in their barbed end affinity, they all similarly increase the actin elongation rate of both leading and trailing barbed ends of fascin-bundled filaments.

**Table 5:** p-values for comparisons of fold change in actin elongation rate with different Ena/VASPs on both leading and trailing filaments of fascin bundles.

| Leading/<br>Trailing<sup>a</sup> | Ena<sub>Tetramer</sub> | VASP | UNC-34 | Ena<sub>Trimer</sub> | Ena<sub>Dimer</sub> |
| --- | --- | --- | --- | --- | --- |
| **Ena<sub>Tetramer</sub>** | 1 / 1 | 0.7 / 0.8 | 0.6 / 0.6 | <u>0.05</u> / 0.2 | <u>0.03</u> / 0.3 |
| **VASP** | 0.7 / 0.8 | 1 / 1 | 0.5 / 0.8 | 0.3 / 0.5 | 0.3 / 0.4 |
| **UNC-34** | 0.6 / 0.6 | 0.5 / 0.8 | 1 / 1 | 0.09 / 0.4 | 0.1 / 0.4 |
| **Ena<sub>Trimer</sub>** | <u>0.05</u> / 0.2 | 0.3 / 0.5 | 0.09 / 0.4 | 1 / 1 | 0.3 / 0.8 |
| **Ena<sub>Dimer</sub>** | <u>0.03</u> / 0.3 | 0.3 / 0.4 | 0.1 / 0.4 | 0.3 / 0.8 | 1 / 1 |

<sup>a</sup> p-values from student's two-tailed t-test with unequal variance between fold change of actin elongation rates when Ena/VASP is bound to the Leading or Trailing barbed end. Underlining shows p-values $\leq 0.05$ .

To test if different Ena/VASP homologs have similarly enhanced processive properties on fascin bundles, two-color TIRFM visualization of 1.5 μM Mg-ATP-actin (15% Oregon Green) was used to quantify the processive run lengths of fluorescently labeled VASP and UNC-34 on fascin bundles (Figure 2E-G, Movie 2). The average residence time of both VASP (1.0 s) and UNC-34 (1.2 s) on single filament barbed ends is ~9-fold shorter than Ena (8.9 s), as expected from lower apparent affinities for barbed ends and previously reported values (Hansen and Mullins, 2010). Yet, like Ena, both VASP and UNC-34 have ~2.5-fold longer processive run lengths on trailing barbed ends of 2-filament bundles (*τ*_VASP_ = 2.6 s, *τ*_UNC-34_ = 2.9 s), with an additional ~1.5-fold increase on trailing barbed ends of 3- or more filament bundles (*τ*_VASP_ = 4.2 s, τ_UNC-34_ = 3.9 s) (Figure 2E–G, Table 1). Therefore, enhanced processivity on fascin-bundled trailing barbed ends is conserved from worms to flies to humans, suggesting that enhanced processivity is important for Ena/VASP’s activity in cells.

### Enhanced elongation and processive run length increases with the number of Ena arms

Wildtype Ena is a tetrameric protein (Kuhnel et al., 2004; Winkelman et al., 2014), with four arms that could facilitate simultaneous associations with a barbed end, neighboring actin filaments, and/or actin monomers for processive elongation. Since we observed that Ena’s average processive run length increases with number of fascin-bundled filaments (Figure 2), we investigated the importance of Ena’s oligomeric state by measuring actin elongation and processive properties of dimeric and trimeric Ena. Dimer and trimer constructs were formed by replacing Ena’s coiled-coil tetramerization domain with a GCN4 dimerization domain (Harbury et al., 1993) or a Foldon trimerization domain (Figure 3A) (Güthe et al., 2004; Papanikolopoulou et al., 2004); and the oligomeric state was verified by gel filtration and multi-angle light scattering (Figure 3 – figure supplement 1A-C). Two-color TIRFM was used to visualize 1.5 μM Mg-ATP-actin (15% Alexa-488 labeled) with SNAP(549)-EnaΔLΔCC-GCN4 (referred to as Ena_Dimer_) or SNAP(549)-EnaΔLΔCC-Foldon (referred to as Ena_Trimer_) on fascin bundles. First, we measured actin elongation rates of Ena-bound leading and trailing barbed ends (Figure 3B, Tables 2,5). While all constructs increase actin’s elongation rate on both leading and trailing filaments, the fold increase is positively correlated with the number of Ena arms. Ena_Tetramer_ has the largest enhancement of actin elongation (2.56-fold leading, 2.1-fold trailing), followed by Ena_Trimer_ (1.74-fold leading, 1.62-fold trailing), and then Ena_Dimer_ (1.45-fold leading, 1.46-fold trailing).

**Figure 3:**
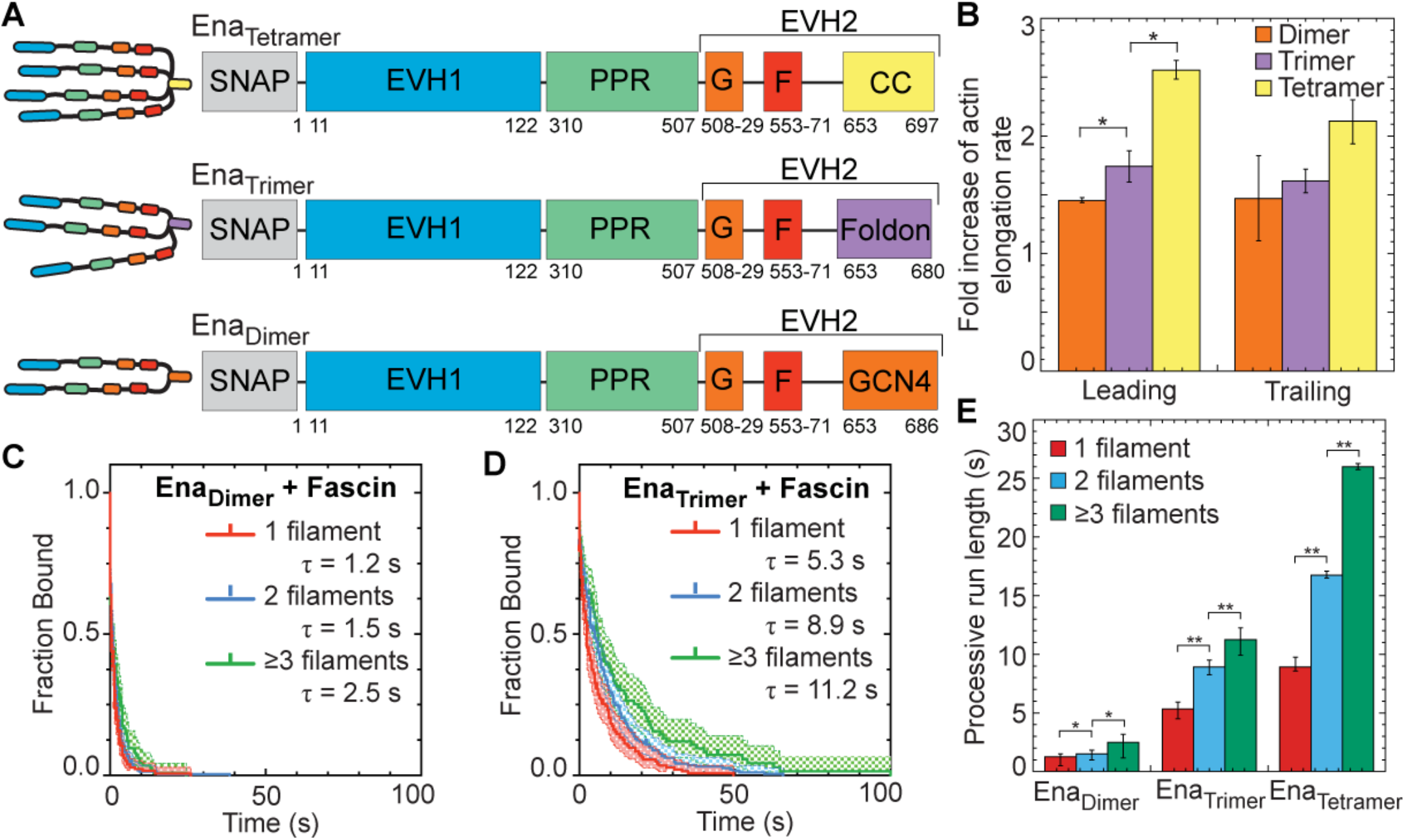
Ena’s processive run length increases with the number of Ena ‘arms’. (A) Cartoon and domain organizations of Ena_Tetramer_, Ena_Trimer_, and Ena_Dimer_. (B-E) Two-color TIRFM visualization of 1.5 μM Mg-ATP-actin (15% Alexa488-actin) with indicated SNAP(549)-Ena construct and 130 nM fascin. (B) Fold increase of barbed end elongation rates of Ena_Dimer_ (orange), Ena_Trimer_ (purple), and Ena_Tetramer_ (yellow). Error bars, SEM. n ≥ 5 barbed ends from at least 2 movies. P values (*≤0.05) (C and D) Kaplan-Meier curves representing average processive run lengths (τ) for (C) 50 pM MBP-SNAP(549)-EnaΔLΔCC-GCN4 or (D) 70 pM MBP-SNAP(549)-EnaΔLΔCC-Foldon with fascin on single filaments (red), or bundles with 2 (blue) and ≥3 (green) filaments. Error bars, 95% CI. n ≥ 93. (E) Average processive run length for increasing number of Ena ‘arms’ on single filaments (red), or fascin bundles with 2 (blue) and ≥3 (green) filaments shown in C and D. Error bars, 95% CI. P values (*<0.05, **<0.0001). Ena_Tetramer_ data in (B) and (E) was also reported in Figure 1K and 2D respectively.

Similar to actin elongation rates, average processive run length is also positively correlated with number of Ena arms (Figure 3C-E). Remarkably, although reduced ~10-fold compared to Ena_Tetramer_, Ena_Dimer_ does remain processively associated with single filament (τ = 1.2 s), 2-filament trailing (τ = 1.5 s), and 3- or more filament trailing (τ = 2.5 s) barbed ends (Figure 3C,E, Movie 3, Table 1). Ena_Trimer_ has intermediate processivity on single filament (τ = 5.3 s), 2-filament trailing (τ = 8.9 s), and 3- or more filament trailing (τ = 11.2 s) barbed ends (Figure 3D,E, Table 1). For each construct, the fluorescence intensity was not correlated with run length (Figure 3 – figure supplement 1D-G), indicating that processive activity is not affected by Ena construct multimerization. Ena_Trimer_’s processive run lengths are similar to the residence time of Ena_Tetramer_ on single filaments but are not comparably enhanced on trailing barbed ends (Figure 3E). Therefore, Ena_Dimer_ is sufficient for processive elongation, Ena_Trimer_ is necessary for longer processive runs on single filaments, but Ena_Tetramer_ is necessary for the longest processive runs on trailing barbed ends of fascin bundles (Figure 3E). Interestingly, the avidity effect of multiple filaments in a fascin bundle is apparent even with fewer arms than the wildtype tetramer. The positive correlation between processive elongation and Ena arms is consistent with a recent study on chimeric human VASP with *Dictyostelium* GAB domains on single actin filaments (Brühmann et al., 2017).

### Tetrameric Ena is more efficient at forming filopodia in *Drosophila* culture cells

Ena_Tetramer_ is significantly better at processive actin filament assembly than either Ena_Dimer_ or Ena_Trimer_, where Ena_Tetramer_ increases the actin elongation rate ~2- to 2.5-fold and remains processively associated with trailing barbed ends of fascin bundles for ~25 sec (Figure 3B,E). To determine whether WT Ena_Tetramer_ is therefore necessary for proper function in cells, we evaluated the ability of Ena oligomerization constructs to facilitate filopodia in ML-DmD16-c3 *Drosophila* culture cells, derived from third instar larval wing discs (Figure 4). We knocked down endogenous Ena with dsRNAi against the 3’UTR and then expressed mCherry-Ena (referred to as mCherry-Ena_Tetramer_), mCherry-EnaΔCC-GCN4 (referred to as mCherry-Ena_Dimer_) or mCherry-EnaΔCC-Foldon (referred to as mCherry-Ena_Trimer_) constructs from a constitutive pIZ plasmid (Figure 4A-C). The activity of the different Ena constructs was determined by quantifying filopodia density, the number of filopodia per perimeter of the cell (Figure 4D). Compared to control cells (0.19 ± 0.06 filopodia/micron), RNAi treated cells without exogenous Ena have a 2.7-fold decrease in filopodia density (0.07 ± 0.03 filopodia/micron). Strikingly, mCherry-Ena_Tetramer_ forms significantly more filopodia (0.24 ± 0.05 filopodia/micron) compared to mCherry-Ena_Trimer_ (0.15 ± 0.05 filopodia/micron) and mCherry-Ena_Dimer_ (0.15 ± 0.04 filopodia/micron). There was no correlation between filopodia density and GFP-actin fluorescence or mCherry fluorescence (Figure 4 – figure supplement 1). Therefore, Ena tetramers facilitate the production of significantly more filopodia than dimer and trimer constructs following knockdown of endogenous Ena.

**Figure 4:**
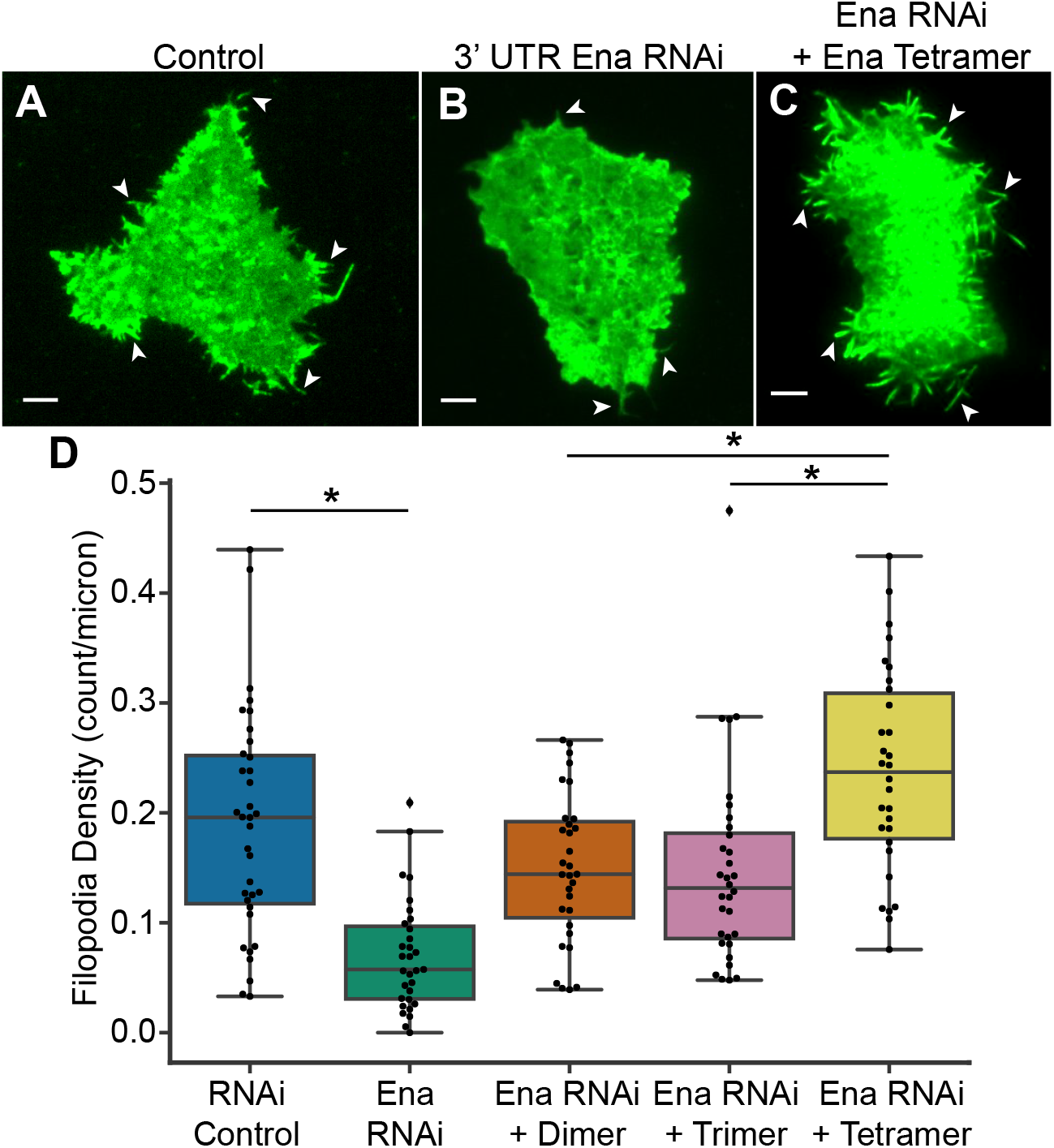
Tetrameric Ena is necessary for proper filopodia density. (A-C) Representative fluorescent micrographs of D16 cells with GFP-actin for (A) Control treatment, (B) Ena 3’ UTR RNAi, and (C) RNAi with transfection of mCherry-Ena_Tetramer_. White arrows indicate representative filopodia. (D) Boxplot of filopodia density, number of filopodia per micron of cell perimeter, for control cells, Ena 3’UTR RNAi, and RNAi transfected with mCherry-EnaΔCC-GCN4 (Ena_Dimer_), mCherry-EnaΔCC-Foldon (Ena_Trimer_), and mCherry-Ena_Tetramer_. n = 3 with at least 10 cells for each experiment. P values (*<0.0005).

### Kinetic model of Ena shows a direct correlation between processivity and both bundle size and Ena oligomerization

We observed that Ena’s processivity depends on the number of filaments in a fascin bundle (Figure 2D) and number of Ena arms (Figure 3E). Therefore, it is likely that the underlying molecular mechanism for Ena’s increased processivity on trailing barbed ends depends on Ena’s ability to simultaneously bind to an elongating barbed end and sides of filaments via its multiple arms (Figure 1B). To investigate this avidity effect, we developed a kinetic model of Ena with varying number of arms, *N*, binding bundles composed of varying number of actin filaments, *n* (Figure 5, Figure 5 – figure supplement).

**Figure 5:**
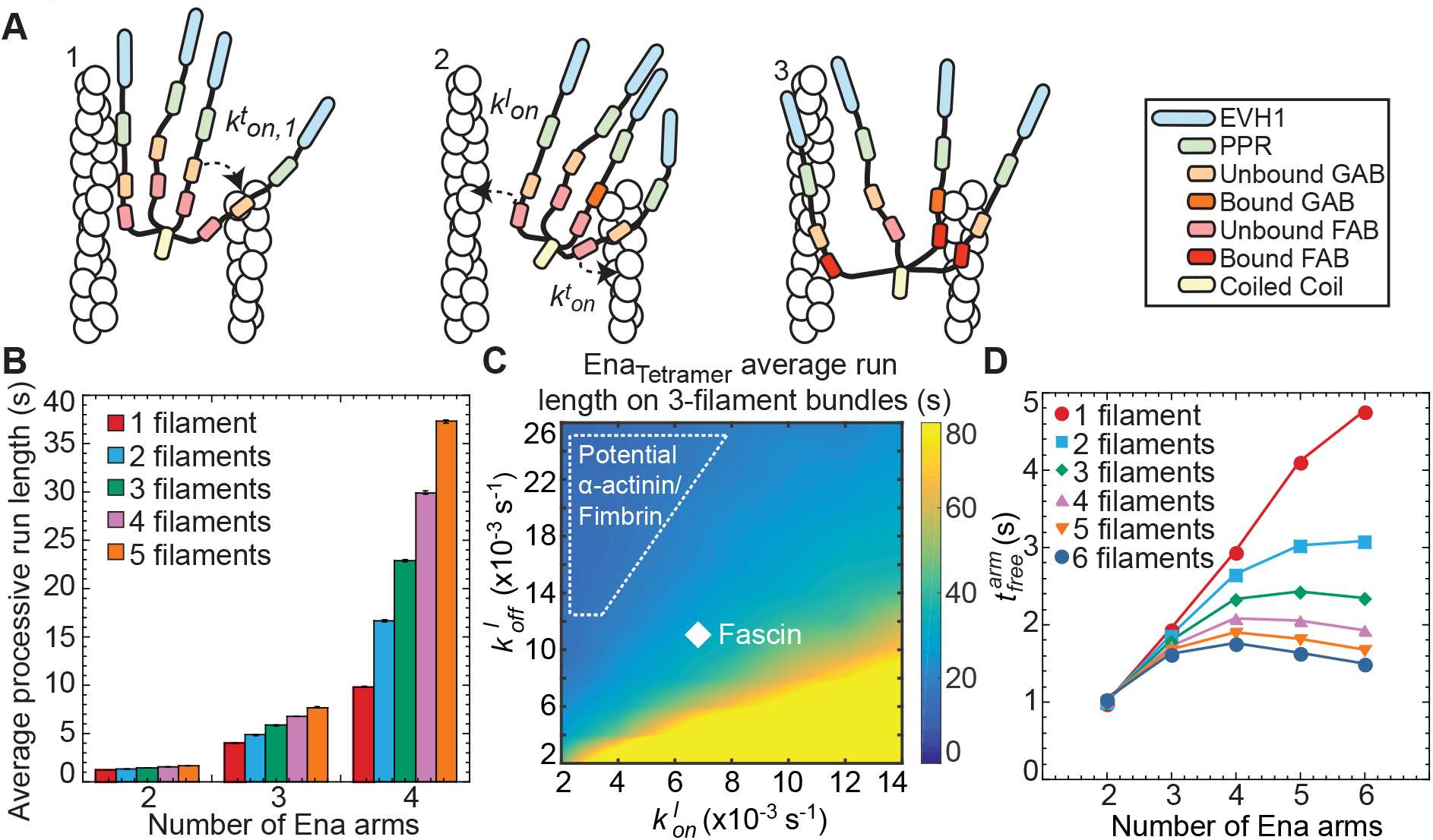
Kinetic model of Ena/VASP on actin bundles shows processivity positively correlates with both number of Ena arms and bundle size. (A) Modelingschematics showing (from left to right). [1] An Ena arm’s GAB domain bindr the trailing barbedend with binding rate 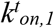. [2] Once the GAB domain is bound, the FAB domain from the other arms binds to sides of either the trailing filament 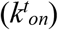 or leading filaments 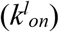. [3] Arms can be bound to the trailing filament, while others bind leading filaments. (B) Bar graph of the average processive run length as a function of number of Ena arms and bundle size. Error bars, SEM. (C) Heat map showing average Ena run length in the case of 3-filament bundles and four Ena arms, with systematic variations of 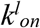 and 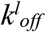. Diamond denotes optimized rates for fascin bundles and region within dotted line shows potential rates for α-actinin and fimbrin. (D) Average time between binding events 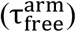 for varying arm number and bundle size.

Our model considers binding and unbinding kinetics of all *N* Ena arms on various binding sites of individual actin filaments in a bundle, which together dictate the kinetics of the Ena “molecule” as a whole (Figure 5A). An Ena arm initially binds to the trailing barbed end with an on rate of 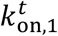 and unbinds with an off rate of 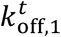 (Figure 5A1). The remaining Ena arms are available to bind and unbind to the side of the trailing filament with a rate 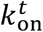 and 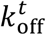 or to the side of other filaments in the bundle with a rate 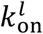 and 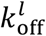 (Figure 5A2-3). A Monte Carlo algorithm was used to integrate rates of binding and unbinding of Ena arms over time as described in the materials and methods. The model parameter 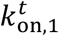 was 0. 007 s^-1^, estimated using the TIRFM measured off rate of 0.109 s^-1^ for Ena, and an equilibrium constant of Ena for the barbed end of 0.8 nM (Winkelman et al., 2014). We therefore considered the local concentration of Ena near the barbed end as 50 pM. The other model parameters were optimized using TIRFM off rates for *N* ∈ (2,3,4) and *n* ∈ (1,2, ≥ 3) (Figure 3E), as described in the materials and methods.

We used the model to characterize Ena’s processive run length at the trailing barbed end. Increasing both the number of filaments in a bundle and the number of Ena arms increases Ena’s processive run length, which strongly supports the avidity hypothesis. The modeling results are also in excellent agreement with the trends observed from our TIRFM data (Figure 5B). Using the model, we tested conditions over a range of both 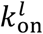 and 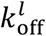 to mimic α-actinin and fimbrin bundles (Figure 1I), where Ena processivity is not enhanced on trailing barbed ends (Figure 5 – figure supplement 1B-F). The model shows a broad regime that results in the same average processive run length on both leading and trailing barbed ends (Figure 5C, dashed region). This indicates that differences between bundlers could be due to diverse association and dissociation rates caused by differences in how CH domain bundlers and fascin bind F-actin.

Finally, we used the model to estimate rates of Ena-mediated filament elongation. While at least one Ena arm associates with the barbed end, its other arms undergo binding and dissociation events. When free, an arm can bind G-actin from solution and transfer it to the barbed end. The elongation rate of the Ena bound filament should be proportional to the average time that individual arms are free. From the model, the average time that individual arms remain unbound while the Ena molecule is in the bound state, 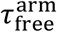, increases with *N*, and decreases with *n* (Figure 5D). This result is consistent with the TIRFM data for the fold increase of actin elongation rate due to Ena on the leading (*n* = 1 in the model) and trailing barbed ends (*n* > 1 in the model) (Figure 1K, 3B).

## DISCUSSION

### Ena’s processivity is enhanced specifically on fascin bundles

Ena/VASP proteins are important processive actin elongation factors that are localized to diverse F-actin networks composed of filaments bundled by different crosslinking proteins, including fascin, fimbrin, and α-actinin. Previously, we found that Ena takes ~3-fold longer processive runs on trailing barbed ends of fascin-bundled F-actin (Winkelman et al., 2014). Here we investigated the mechanism and conservation of Ena/VASP’s processivity at the barbed end of single filaments and filaments bundled by different crosslinking proteins, as well as the physiological relevance of Ena/VASP tetramerization.

We found that although fly Ena’s processivity is enhanced ~3-fold on trailing barbed ends in fascin bundles, there is no processivity enhancement on trailing barbed ends of α-actinin or fimbrin bundles (Figure 1I). Fimbrin and α-actinin use two CH domains to bundle F-actin, whereas fascin uses β-trefoil domains. Though the exact mechanism for Ena’s specificity for fascin bundles remains unclear, we suggest several hypotheses. First, fascin could hold the trailing filament in a specific register with respect to the leading filament, allowing for easier Ena/VASP binding. Second, fascin’s strong cooperativity (Winkelman et al., 2016; Yamakita et al., 1996) could promote more rapid bundling, thereby promoting longer processive runs by keeping trailing barbed ends closer to sides of leading filaments. Third, it is also possible that Ena weakly associates with fascin, although no interaction has yet been detected. If Ena does associate with fascin, it would need to be carefully tuned because a strong interaction could pull Ena from the barbed end. Fourth, our kinetic model revealed a broad region of Ena binding kinetics to sides of bundled filaments 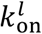 and 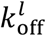 that could explain Ena’s lack of enhanced processivity on fimbrin and α-actinin bundles (Figure 5B). It is possible that these rates are affected by competition between Ena and the CH domain bundling proteins for similar binding sites on actin filaments. Further studies of how fascin forms F-actin networks differently than α-actinin and fimbrin will be required to fully elucidate the underlying molecular mechanism. However, this important observation reveals for the first time that bundling proteins and the F-actin networks they form can differentially regulate the activity of processive actin assembly factors, thereby providing a mechanism to allow Ena/VASP proteins to facilitate the assembly of diverse bundled networks with different dynamics in cells. Understanding how different bundling proteins associate with and help form specific F-actin networks in cells will therefore be of critical importance.

### The mechanism of tetrameric Ena acting on fascin bundles for filopodia formation

Given that Ena localizes to filopodia with fascin, lamellipodia with fimbrin and stress fibers with α-actinin, sensitivity to diverse bundles could play an important role in regulating Ena activity in cells. Filopodia are unique amongst these networks with long, straight filaments that emerge from a network capped by capping proteins. Lamellipodia have short, branched filaments and stress fibers are contractile, bipolar networks. Thus, filopodia are the ideal network for enhanced Ena/VASP processivity facilitating elongation of longer filaments that requires stronger competition against capping protein to form a protrusive network. The increased residence time on trailing barbed ends could play a critical role in a feedback mechanism between Ena and fascin in emerging filopodia (Winkelman et al., 2014). Ena/VASP-associated barbed ends elongate faster, assembling longer actin filaments that contain more fascin binding sites, which subsequently enhance Ena/VASP’s processivity. Trailing barbed ends that have longer Ena processive runs can catch up to the leading barbed end, allowing all filaments to reach the same length and resulting in mature filopodia with uniform thickness and aligned barbed ends.

### Avidity promotes enhanced Ena processivity on fascin bundles

We hypothesize that avidity between multiple actin filaments in a fascin bundle and multiple Ena arms promotes the formation of long filopodia filaments. We investigated the avidity effect by testing how the number of filaments in a fascin bundle and number of Ena arms affects Ena’s processive run length. Our results strongly indicate that avidity plays a major role, as there is a ~2-fold increase in Ena’s residence time on trailing barbed ends in 2-filament bundles and an additional ~1.5-fold increase on bundles with 3 or more filament compared to single filament barbed ends (Figure 2B-D). Similarly, the residence time of both VASP and UNC-34 is longer on trailing barbed ends and is correlated with number of actin filaments in a fascin bundle (Figure 2E-G). Furthermore, the residence time of Ena_Trimer_ and Ena_Tetramer_ is ~4.5- and ~10-fold longer than Ena_Dimer_ on fascin bundles with 3 or more filaments (Figure 3C-E). A recent study measuring processive elongation using chimeric human VASP with *Dictyostelium* FAB domains on single filaments (Brühmann et al., 2017) supports our conclusions that enhanced elongation and processive run length are positively correlated with the number of Ena arms. Observing this positive correlation under more ‘physiological conditions’, a construct using Ena’s unmodified EVH2 domains and on fascin bundles, indicates that these properties are relevant for Ena’s activity in cells and specifically for filopodia.

We further tested the avidity hypothesis by developing a kinetic model that incorporates Ena with differing number of arms binding to single or multiple filaments (Figure 5). Previous models have focused exclusively on modeling the kinetics of Ena/VASP-mediated barbed end elongation of single actin filaments (Breitsprecher et al., 2011; Brühmann et al., 2017; Hansen and Mullins, 2010). VASP-mediated single filament elongation rates were shown to increase linearly with the number of VASP arms in solution as predicted by the model (Breitsprecher et al., 2011). However, this model overlooks the binding kinetics of arms that are not associated with the barbed end. Hence, we developed a kinetic model that explicitly incorporates the binding and unbinding rates of each Ena arm on multiple filaments (Figure 5A). After an Ena arm binds to the barbed end 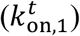, the remaining arm(s) are free to bind to the side of the leading filament(s) 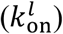 or the trailing filament 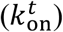. We quantified the processive run length for various numbers of bundled filaments and Ena arms.

The model demonstrates that the avidity effect of Ena emerges from an effective increase in local concentration of F-actin that allows for more FAB binding sites and from multiple Ena arms with available FAB domains. The avidity effect results in longer residence times near the trailing barbed end. Importantly, if an arm dissociates from the trailing barbed end, Ena will continue to processively elongate the barbed end and not diffuse away given that other arms’ FAB domains are associated with nearby actin filaments. Furthermore, our model that includes multiple arms binding to multiple actin filaments still has a linear correlation of elongation rates with number of Ena arms on single filaments (Figure 5D), as predicted by a previous model (Brühmann et al., 2017). The 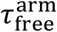 is linear with respect to increasing additional Ena arms on single filaments, but with increasing number of filaments there are diminishing returns by adding more Ena arms. 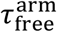 peaks at a tetramer on larger bundles, which gives an additional argument of why a tetramer of Ena/VASP is evolutionarily preferred. We also observe that an Ena tetramer is more efficient at forming filopodia in *Drosophila* culture cells compared to dimer and trimer constructs (Figure 4). Since the tetramer has increased residence time on trailing barbed ends and increases actin’s elongation rate above the dimer and trimer, this suggests that the tetramer is necessary for proper actin elongation rates and competition with capping protein to allow for the formation of the correct number of filopodia.

## MATERIALS AND METHODS

### Total internal reflection fluorescence microscopy (TIRFM)

TIRFM images were collected at 250ms-1s intervals with a cellTIRF 4Line system (Olympus, Center Valley, PA) fitted to an Olympus IX-71 microscope with through-the-objective TIRF illumination and an iXon EMCCD camera (Andor Technology, Belfast, UK). Mg-ATP-actin (15% Oregon Green or Alexa 488 labeled) was mixed with polymerization TIRF buffer [10 mM imidazole (pH 7.0), 50 mM KCl, 1 mM MgCl_2_, 1 mM EGTA, 50 mM DTT, 0.2 mM ATP, 50 μM CaCl_2_, 15 mM glucose, 20 μg/mL catalase, 100 μg/mL glucose oxidase, and 0.5% (400 centipoise) methylcellulose] to induce F-actin assembly and any additional actin binding proteins. This mixture was transferred to a flow cell for imaging at room temperature. For two color TIRFM, we cyclically imaged labeled actin (1 frame, 488 nm excitation for 50ms) and SNAP(549)-Ena/VASP (1 frame, 561 nm excitation for 50ms) (Winkelman et al., 2014).

### D16 cell culture

ML-DmD16-c3 (DGRC) cells were cultured in Schneider’s Media with 10% Fetal Bovine Serum (Gibco, Waltham, MA), Anti-Anti (Gibco, Waltham, MA), and 10 μg/mL recombinant human insulin (Gibco, Waltham, MA), transfected with FugeneHD (Promega, Madison, WI), and imaged on extracellular matrix (ECM) coated glass-bottom dishes after 48-72 hr. ECM was harvested from ML-DmD17-c3 (DGRC, Bloomington, IN) (Currie and Rogers, 2011). All imaging was performed on a total internal reflection fluorescence (TIRF) system mounted on an inverted microscope (Ti-E, Nikon, Tokyo, Japan) using a 100X/1.49NA oil immersion TIRF objective driven by Nikon Elements software unless noted otherwise. Images were captured using an Orca-Flash 4.0 (Hamamatsu, Hamamatsu, Japan) and were processed for brightness and contrast using ImageJ (Schneider et al., 2012) analysis. We quantified >30 cells using CellGeo (Tsygankov et al., 2014). Filopodia were quantified with the criteria of >0.78 μm long and <0.91 μm wide.

### Plasmid Construction

Enabled (Ena) constructs were prepared by removing the 6x-His tag from the C-terminus of previously described Ena constructs [MBP-SNAP-EnaΔL or MBP-EnaΔL] (Winkelman et al., 2014) and insertion into a MBP containing plasmid (pet21A) by standard restriction digest and infusion (Clontech, Mountain View, CA) following PCR amplification (iProof; Bio-Rad, Hercules, California). Ena_Dimer_ and Ena_Trimer_ constructs were prepared by removing the coiled-coil domain and adding a Foldon domain (Güthe et al., 2004; Papanikolopoulou et al., 2004) [MBP-SNAP-EnaΔLΔCC-Foldon] or GCN4 domain (Harbury et al., 1993) [MBP-SNAP-EnaΔLΔCC-GCN4] from MBP-SNAP-EnaΔL. UNC-34 was cloned from worm cDNA and inserted into a pet21A vector with MBP-SNAP (New England Biolabs, Ipswich, MA) at XmaI/PacI sites, while also including a flexible linker (GGSGGS) in the forward primer sequence of SNAP constructs. Singed and VASP constructs were cloned from fly and human cDNA libraries, respectively. VASP was inserted into a MBP-SNAP and SNAP containing vector while Singed was inserted into a pGEX KT Ext plasmid containing GST with a Thrombin cleavage site at XbaI/XhoI sites. Plasmids for transfection of mCherry-EnaΔCC-GCN4 and mCherry-EnaΔCC-Foldon were cloned into a pIZ-mCherry-Ena (Bilancia et al., 2014) construct using infusion (Clontech, Mountain View, CA). The RNAi was designed using Primer3Plus (Untergasser et al., 2012) targeting the 3’ UTR of *enabled* using forward primer 5’ TAATACGACTCACTATAGGGAGACCACGTGATGGCATGTGCATAGGC 3’ and reverse primer 5’ TAATACGACTCACTATAGGGAGACCACTGCTGAAGACTTGCTGGTTC 3’. The 3’UTR was extracted from w1118 strain fly genome and the DNA region of interest was isolated by PCR amplification and placed in a bluescript SK vector. dsDNA was produced using PCR amplification and dsRNA was produced from the resulting dsDNA using MEGAscript T7 Transcription kit (Invitrogen, Waltham, MA).

### Protein Expression and Purification

Recombinant Ena/VASP proteins were purified by expressing in Escherichia coli strain BL21-Codon Plus (DE3)-RP (Agilent Technologies, Santa Clara, CA) with 0.25 mM isopropyl β-D-1-thiogalactopyranoside for 16 h at 16 °C. Cells were lysed with an Emulsi-Flex-C3 (Avestin, Ottawa, Canada) in extraction buffer [20 mM TRIS-HCl (pH 8.0), 200 mM NaCl, 10% glycerol, 0.1 mM DTT] with 0.5 μM PMSF and cOmplete, EDTA-free Protease Inhibitor Cocktail (Roche, Basel, Switzerland) and were clarified. The extract was incubated for 1 h at 4 °C with amylose resin (New England Biolabs, Ipswich, MA) and was washed with extraction buffer, then Ena/VASP was batch eluted with elution buffer [20 mM TRIS-HCl (pH 8.0), 200 mM NaCl, 10% glycerol, 0.1 mM DTT, 40 mM maltose] Ena/VASP was incubated overnight with and without 1 uM TEV protease to cleave MBP and filtered on an Superdex 200 10/300 GL or Superose 6 Increase 10/300 GL column (GE Healthcare, Little Chalfont, UK) where they eluted as stable oligomers. Ena/VASP constructs were dialyzed against SNAP buffer [20 mM Hepes (pH 7.4), 200 mM KCl, 0.01% NaN_3_, and 10% Glycerol, and 0.1 mM DTT]. SEC-MALS was performed using DAWN HELEOS II and Optilab T-rEX (Wyatt Technology, Goleta, CA) with a Superdex 200 Increase 10/300 GL column and Akta FPLC (GE Healthcare, Little Chalfont, UK). SEC-MALS data was analyzed using Astra 6.0 (Wyatt Technology, Goleta, CA). SNAP-tagged proteins were labeled with BG-549 (New England Biolabs, Ipswich, MA) following the manufacturers’ protocols. Concentrations of SNAP-tagged proteins and the degree of labeling were determined by densitometry of Coomassie stained bands on SDS/PAGE gels compared with standards. Ena/VASP was flash-frozen in liquid nitrogen and stored at −80 °C. N-terminal SNAP and MBP tags did not affect Ena/VASP’s activity. Actin was purified from rabbit skeletal muscle acetone powder (Pel-Freez, Rogers, AR) or self-prepared chicken skeletal muscle acetone powder by a cycle of polymerization and depolymerization and gel filtration (Spudich and Watt, 1971). Gel-filtered actin was labeled with Oregon green (Kuhn and Pollard, 2005) or Alexa 488. Human fascin, human α-actinin IV, and S. pombe fimbrin were expressed in bacteria and purified as described (Li et al., 2016; Skau and Kovar, 2010; Vignjevic et al., 2003). Singed was purified in the same manner as previously reported for human fascin (Vignjevic et al., 2003).

### Glass Preparation

Microscope slides and coverslips (#1.5; Fisher Scientific, Waltham, MA) were washed for 30 min with acetone and for 10 min with 95% ethanol, were sonicated for 2 h with Helmanex III detergent (Hellma Analytics, Müllheim, Germany), incubated for 2 h with piranha solution (66.6% H_2_SO_4_, 33.3% H_2_O_2_), washed with deionized water, and dried. Glass then was incubated for 18 h with 1 mg/mL mPeg-Silane (5,000 MW) in 95% ethanol, pH 2.0. Parallel strips of double-sided tape were placed on the coverslip to create multiple flow chambers (Zimmermann et al., 2016).

### Calculation of Residence Time and Elongation Rates

To calculate Ena/VASP’s residence time on barbed ends, SNAP(549)-Ena/VASP fluorescent spots associated with the barbed end were manually tracked using MTrackJ (Meijering et al., 2012) in ImageJ. Spots that did not move were not scored, because they were assumed to be adsorbed to the glass. Events that contained joined barbed ends with no clear leading or trailing barbed bend were not included in the average lifetime calculation. Residence times for single SNAP-549-Ena(ΔL) tetramers were determined by fitting a Kaplan-Meier (Kaplan and Meier, 1958) survival curve with a single exponential equation, f(x) = x0 * exp^(−x/T1) to calculate the average lifetime. Kaplan Meier survival curves were used to account for processive runs that started before imaging began or ends after imaging terminated. Log rank statistical significance tests were done using Prism 7 (GraphPad Software, San Diego, CA). Barbed end elongation rates were calculated by measuring filament lengths over time with ImageJ software. Multiple filament lengths were plotted over time and the distribution was fit with a linear equation using KaleidaGraph 4.5 (Synergy Software, Reading, PA). To calculate the number of filaments in a bundle the TIRFM movie was used to follow the history of the filaments. This could most accurately differentiate between two-filament bundles and three or more filament bundles. Due to photobleaching of the filaments over time the actin fluorescence was not used to determine the number of filaments within the bundle.

### Fluorescence Spectroscopy

Bulk actin assembly was measured from the fluorescence of pyrene-actin with a Safire2 or Infinite M200 Pro (Tecan Systems, Inc., Männedorf, Switzerland) fluorescent plate reader (Neidt et al., 2008). Briefly, unlabeled Mg-ATP-actin was preassembled into seeds for 1 hour by adding 50 mM KCl, 1 mM MgCl_2_, 1 mM EGTA, 10 mM imidazole, pH 7.0. The assay measures the elongation rate of actin by addition of 20% pyrene-labeled Mg-ATP-actin monomers and actin binding proteins to be assayed. Final protein concentrations are indicated in the figure legends.

### Development of kinetic model

In order to test, mechanistically, the hypothesis that avidity of Ena binding multiple actin filaments with multiple arms determines an increase in time spent at the trailing barbed end for fascin-crosslinked bundles, we developed a computational model. The model is based on a kinetic Monte Carlo algorithm that at each time step evaluates binding and unbinding probabilities of each Ena arm for each filament and, accordingly, changes the arm “state”. The kinetic Monte Carlo scheme is chosen because it can, in principle, give the exact evolution of the system, in terms of bound and unbound states of each Ena arm over time, thus providing a strong approximation of the sequence of events given individual Ena arm’s binding and unbinding rates, with respect to individual filaments. The kinetic model used in this work consisted of the following elementary reactions:

1. Initial binding of an arm of Ena to the barbed end of the trailing filament with a rate of 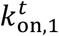.
2. At every subsequent step, binding and unbinding of:

a. an arm of Ena to the barbed end of the trailing filament, with rates 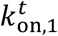 and 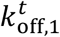
b. up to two other arms of Ena to the side of the trailing filament, with rates 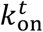 and 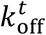
c. additional arms of Ena beyond three to the side of the trailing filament, with rates 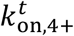 and 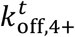
d. other arms of Ena to the sides of other filaments in the bundle, with rates 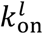 and 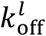

 In summary, once any arm is bound to the barbed end of the trailing filament, the Ena “molecule” is considered to be in the bound state. The Ena molecule unbinds only when none of its arms are bound to any of the filaments in the bundle. Thus, after initiation of the bound state for an Ena molecule, the arm bound to the barbed end can unbind and bind multiple times before the molecule unbinds.

The model was made efficient by only simulating events involving binding and unbinding of the Ena molecule to the barbed end of a trailing filament. Further, we did not intend to calculate the binding rate of the Ena molecule using the model, and instead optimized the model parameters based on TIRFM data (see below) for calculating the unbinding rates of Ena molecules, and predicting the kinetics of individual Ena arms while the molecule was bound. This gave rise to the following possible scenarios while the Ena molecule is in the bound state.

1. Only one arm is bound to either

a. the barbed end
b. the side of the trailing filament
c. the side of another filament in the bundle
2. Two or more arms are bound

a. one to the barbed end, others to the side of the same filament
b. one to the barbed end, others to the side of another filament
c. one to the barbed end, others to the sides of the same and other filament(s)
d. some to the side of the same filament and the remaining to the side of another filament
e. all to the side of the same filament
f. all to the side of another filament

### Model parameters

Since Ena is a homotetramer, all arms in this work are structurally identical to each other. Hence, not all of the eight kinetic rate constants 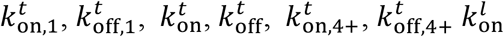 and 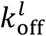 in the model (Figure 5A) are independent. We set 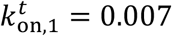, estimated using the TIRFM measured off rate of 0.109 s^-1^ for Ena, and an equilibrium constant of Ena for the barbed end of 0.8 nM (Winkelman et al., 2014). The model assumes that binding rates of the rest of the arms to sides of filaments are identical 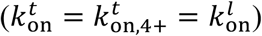, consistent with the idea that avidity results from binding and unbinding of multiple Ena arms to multiple filaments, rather than from different kinetics of individual arms. The corresponding unbinding rates were, however, assumed to be different owing to the following reasons. An arm bound to the barbed end interacts with the barbed end of the filament through its GAB domain and potentially its FAB domain, while an arm bound to the side of a filament interacts only through its FAB domain. Thus, 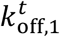 is considered an independent parameter. The number of FAB domain binding sites available on the trailing filament can be assumed to be less than those on leading filaments since it is the shortest filament in the bundle. Thus, 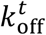 and 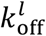 are a priori considered to be distinct parameters. Our TIRFM data (Figure 3B) suggests that the fold increase in processive run length between a trimer and a tetramer binding to a single filament is smaller than the fold increase between a dimer and a trimer. Thus, the fourth arm binding to the same filament is assumed to have different unbinding kinetics represented using the rates 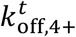. This translates to having an upper limit on the number of arms that can simultaneously bind to a given filament.

### Optimization procedure for parameter estimation

With the above assumptions, the number of undetermined parameters to be estimated reduces to five: 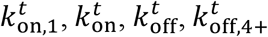 and 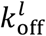. These parameters were estimated using all 9 data points for the processive run length data in Figure 3B using the Levenberg-Marquardt algorithm implemented in the MATLAB^®^ function “fsolve”. Let *τ*(*n, N*) represent the processive run length of Ena with *N* arms on the trailing barbed end of a bundle consisting of *n* actin filaments. The rate ratio vector *y* is defined as

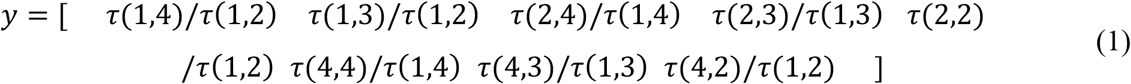

the error was then defined as

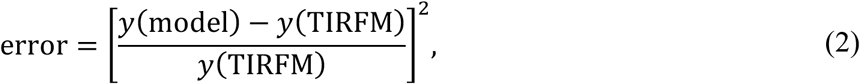

and minimized iteratively using the five undetermined parameters. For each iteration, the kinetic model was solved for each pair of *N* ∈ (2,3,4) and *n* ∈ (1,2,4) and the corresponding average processive run length (defined below) was calculated. The TIRFM data for *n* ≥ 3 in Figure 3E, corresponding to three or more filaments in the bundle, was considered to be equivalent to *n* = 4 in the model, consistent with our observation that most bundles in the TIRF data fell between 3 and 5 filaments for an average ‘large’ bundle.

For computational efficiency, we adopted a two-step strategy to obtain the optimum set of parameters. In the first step, we performed error minimization using 50 distinct initial guesses for the parameters and chose six optimized parameter sets with the lowest errors. In the second step, we performed error minimization using 100 sets of initial guesses, each perturbed within ±10% of the average of these six sets from the first step. The parameter set with the least error was chosen as the final set (Table 6). A comparison of the rate ratio vectors from the model with corresponding data from TIRFM is shown in Table 7. The optimized parameter set was found to predict rate ratios in good agreement with the corresponding ratios from TIRFM data (Figure 3E).

**Table 6:** Final set of rate constants in the kinetic model.

| $k_{\text{on},1}^t$ | $k_{\text{on}}^t = k_{\text{on},4+}^t = k_{\text{on}}^l$ | $k_{\text{off},1}^t$ | $k_{\text{off}}^t$ | $k_{\text{off},4+}^t$ | $k_{\text{off}}^l$ |
| --- | --- | --- | --- | --- | --- |
| 0.007 | 0.0122 | 0.1488 | 0.0049 | 0.0055 | 0.0195 |

**Table 7:** Comparison of processive run length ratios defined in Equation (S1) from the model and from TIRFM data.

| Run length ratios | $\frac{\tau(1,4)}{\tau(1,2)}$ | $\frac{\tau(1,3)}{\tau(1,2)}$ | $\frac{\tau(2,4)}{\tau(1,4)}$ | $\frac{\tau(2,3)}{\tau(1,3)}$ | $\frac{\tau(2,2)}{\tau(1,2)}$ | $\frac{\tau(4,4)}{\tau(1,4)}$ | $\frac{\tau(4,3)}{\tau(1,3)}$ | $\frac{\tau(4,2)}{\tau(1,2)}$ |
| --- | --- | --- | --- | --- | --- | --- | --- | --- |
| **y(model)** | 8.3240 | 3.3356 | 1.6951 | 1.2305 | 1.0902 | 2.9064 | 1.7222 | 1.2969 |
| **y(TIRFM)** | 7.5727 | 4.4074 | 1.8333 | 1.6875 | 1.2489 | 2.8947 | 2.1236 | 2.0825 |

### Algorithm

Using the values of reaction rates provided in Table 6, the system evolved using a Monte Carlo algorithm with a constant time-step implemented in MATLAB^®^. The states of the arm binding the barbed end and the other arms binding sides of filaments was stored along corresponding columns in a *state* array, with an entry of 0 representing an unbound state and 1 representing a bound state. Each row in the array corresponded to a simulation step. The identity of the filament in the bundle that each Ena arm bound to was stored in a separate *filamentid* array, with filament identities ranging from 1 to *n*.

The simulation was initialized with all arms of Ena in the unbound state. At each timestep *t* + *dt*, a reaction move (either binding or unbinding) and the corresponding rate constants were selected depending on the previous state of the system at timestep *t* (Figure 5 – figure supplement 1A). For example, if the barbed end was bound at timestep *t*, the unbinding reaction with the rate constant of 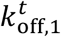 was selected at timestep *t* + *dt. N* random numbers were generated, one corresponding to each arm, and compared with the rate constant of the selected reaction move. The move was accepted if the random number was less than the corresponding rate constant times *dt*, and the entries *state* and *filamentid* arrays were updated accordingly.

### Model verification and predictions

The quantities in the model with units of timesteps were converted to real time in seconds by multiplying with a single factor of 5.4374 × 10^-3^ that accounted for the “timescale” and was chosen to exactly match the processive run length for a dimer on a single filament between the model (defined below) and the TIRFM data (leftmost red bar in Figure 3E). For computational efficiency, we used *dt* = 0.1 s.

Assuming that any difference in fascin, alpha-actinin and fimbrin bundles due to spacing between filaments or different interactions should be reflected in the binding and unbinding kinetics, we systematically varied binding/unbinding rates 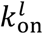 and 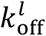 from 0.002 to 0.026 s^-1^, keeping other model parameters fixed (Figure 5 – figure supplement 1B-F). For a single filament, the processive run length of an Ena tetramer is independent from 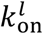 and 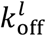 as expected (Figure 5 – figure supplement 1B). With more than one filament, the processive run length increases with 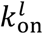, for increasing values of 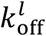 below ~0.010 s^-1^ (Figure 5 – figure supplement 1C) or below ~0.026 s^-1^(Figure 5 – figure supplement 1D). We also systematically changed 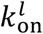 and 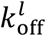 using dimers and trimers on 3-filament bundles. Similar to the Ena tetramer, the Ena trimer and dimer showed processive run lengths that increased with 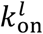 for values of 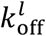 below ~0.014 s^-1^ (Figure 5 – figure supplement 1E-F). In the tested range of values for 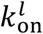 and 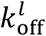, the maximum run time with dimers is 3 s (Figure 5 – figure supplement 1E) and trimers is 20 s (Figure 5 – figure supplement 1F). Our results show that the processive run length is determined by an interplay between the numbers of arms and filaments, and cross-linker effects on binding rates to sides of leading filaments.

### Defining processive run length (*τ*) and free arm time 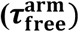

#### Processive run length (*τ*)

The Ena molecule binding was considered the beginning of a processive run event (*τ*_start_, Figure 5 – figure supplement 1A) and unbinding of the Ena molecule (*τ*_end_, marked in Figure 5 – figure supplement 1A) denoted the end of a processive run event. The processive run length τ was calculated by averaging the difference (*τ*_start_ – *τ*_end_) across all processive run events observed across 56 independent simulation runs, each consisting of a total of 2 × 10^6^ timesteps (equivalent to ~10000 seconds).

For the final data in Figure 5B, the total number of processive run events used for averaging varied depending on the number of Ena arms and number of filaments in the bundle. Based on the range in our TIRFM data (Figure 3E), the number of events were in the range of ~1.6 × 10^5^ for (*N* = 4, *n* = 4) and 6.8 × 10^5^ for (*N* = 2, *n* = 1). The least number of events used in obtaining data in Figure 5, ~4.6 × 10^3^, corresponded to (*N* = 6, *n* = 6).

#### Free arm time 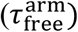

During each processive run event in the model, individual Ena arms bind to and unbind from filaments independently, but according to their specific rates. The average time between consecutive binding events of an average arm was calculated and denoted as 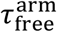 . A free Ena arm is available to recruit G-actin from the solution and transfer it to the barbed end with an effective rate that should be independent of the number of filaments in the bundle. Further, since each arm is identical, the effective rate should also be independent of the identity of the arm. It should be noted that in the model Ena arms do not have an identity associated with them and are only used as proxies to obtain statistics related to occupied versus unoccupied states of the barbed end and the sides of filaments. A rapid exchange of an Ena arm bound to the barbed end with an arm bound to the side of a filament is possible but not explicitly accounted for in the model. Thus, though the kinetic model does not explicitly consider filament elongation, 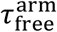 is assumed to be approximately proportional to the elongation rate through this implicit effective rate at the resolution of the model.

## FIGURES AND FIGURE LEGENDS

**Figure 2 – figure supplement 1.**
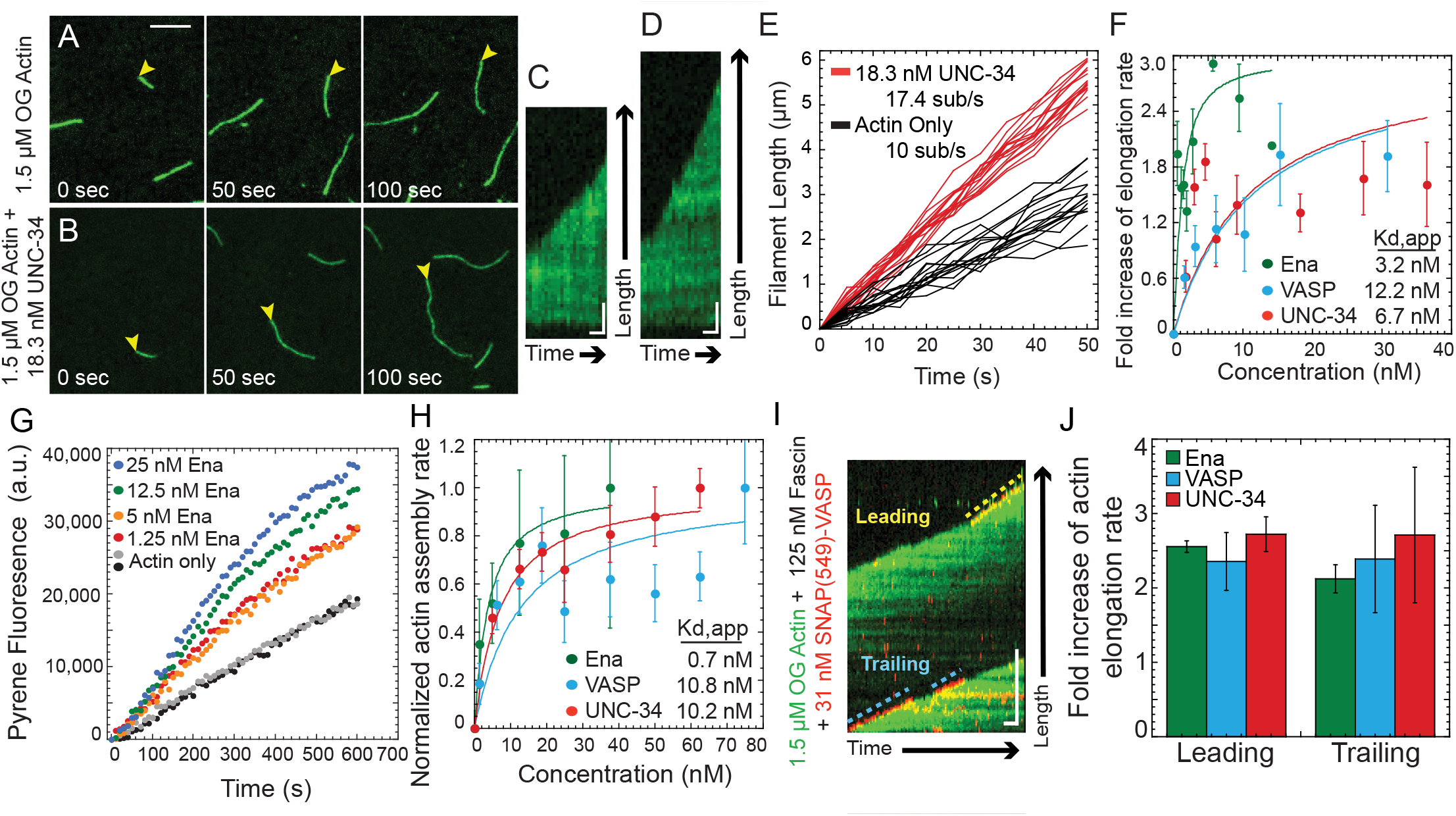
Ena/VASP homologs have generally conserved processive actin elongation properties. (A-F) Single-color TIRFM of the spontaneous assembly of 1.5 μM Mg-ATP-actin (15% Oregon Green) with worm UNC-34, fly Ena, and human VASP. (A and B) Time-lapse micrographs (scale bar, 5 μm), and (C and D) corresponding kymographs (scale bar, 1μm; time bar, 5s) for (A and C) actin alone or with (B and D) 18 nM UNC-34. Yellow arrowheads indicate barbed ends. (E) Length of individual filaments over time for actin only (black) and UNC-34 (red). (F) Fold increase in elongation rate over increasing concentration of Ena (green), VASP (blue), and UNC-34 (red). Curve fits revealed the indicated apparent dissociation constants (Kd, app) of Ena/VASP for the barbed end. n ≥ 5 filaments for at least 2 movies. (G and H) Seeded assembly: addition of 0.5 μM Mg-ATP-actin monomers (20% pyrene-labeled) to the barbed end of 0.5 μM preassembled actin filaments. (G) Time course of seeded assembly alone (black and gray) or with a range of Ena concentrations. (H) Dependence of the initial barbed end assembly rate on Ena/VASP concentration. Curve fits revealed the indicated apparent dissociation constants (Kd, app) of Ena/VASP for the barbed end. Error, SEM. n ≥ 3. (I and J) Two-color TIRFM visualization of 1.5 μM Mg-ATP-actin (15% Oregon green-actin) with 25 pM SNAP(549)-VASP, 18 pM SNAP(549)-UNC-34 or 15 pM SNAP(549)-Ena, and unlabeled 130 nM human fascin. (I) Kymograph of leading and trailing barbed ends of a Fascin bundle with SNAP(549)-VASP (red) and OG actin (green). Dashed blue (trailing) and yellow (leading) lines indicate bound VASP. Scale bar, 5 μm. Time bar, 10 s. (J) Average elongation rate of leading and trailing filament barbed ends on fascin bundles with actin alone, Ena, VASP, or UNC-34. Error bars, SEM. n ≥ 5 filaments for at least 2 movies. Elongation rate for Ena in (J) also shown in Figure 1K.

**Figure 3 – figure supplement 1.**
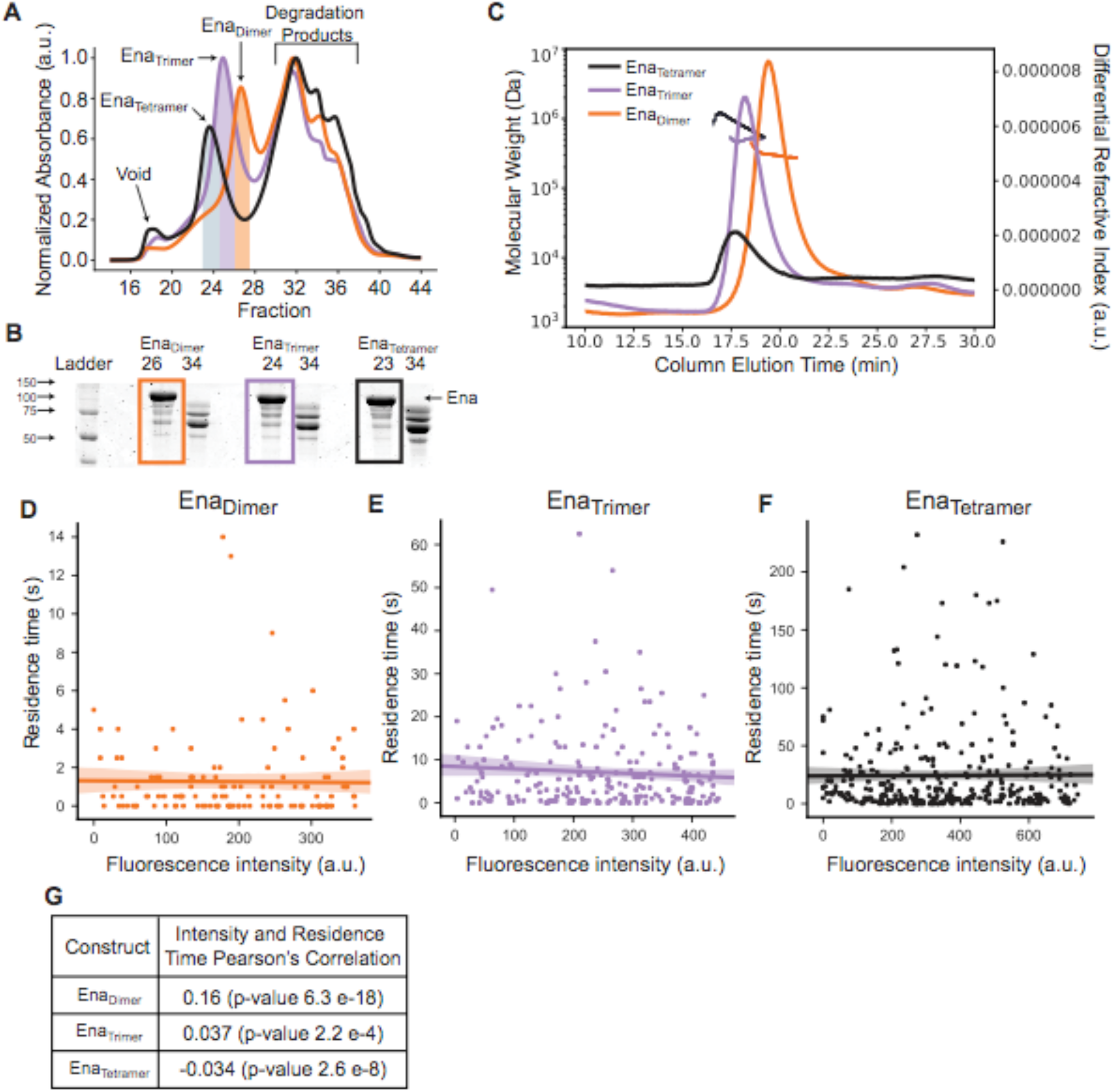
Ena constructs have the predicted oligomerization state. (A) UV traces for Ena_Dimer_ (orange), Ena_Trimer_ (purple), and Ena_Tetramer_ (black) from size exclusion gel filtration using a Sepharose 6 Increase column. Peaks are labeled and fractions collected are shaded. (B) The 12.5% SDS-PAGE of fractions from A. Fractions showing each construct are boxed. (C) Size exclusion chromatography followed by multi-angle light scattering (SEC-MALS) was used to determine the relative size of Ena_Dimer_ (orange), Ena_Trimer_ (purple), and Ena_Tetramer_ (black). (D) Values for the Pearson’s correlation of the Ena construct fluorescence intensity and its residence time for all movies analyzed. There is no correlation between an Ena construct’s intensity and its bound lifetime. Dependence of (E) Ena_Dimer_, (F) Ena_Trimer_, or (G) Ena_Tetramer_ processive run length on its respective fluorescence intensity for an individual movie. Linear correlation fit shown with 95% CI shaded.

**Figure 4 – figure supplement 1.**
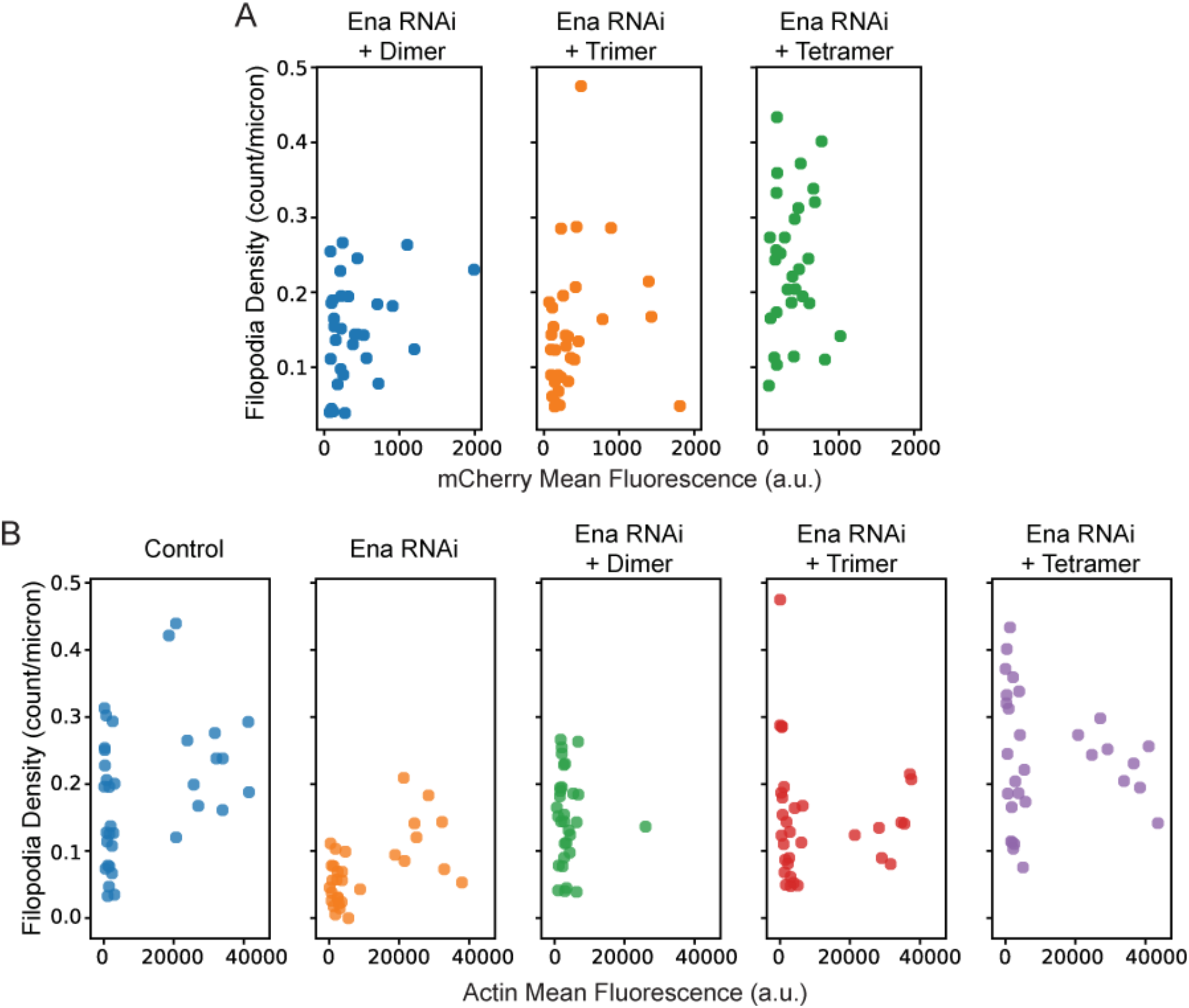
D16 culture cell expression is independent of construct. (A) Western blot showing expression levels of transfected mCherry-Ena_Dimer_, mCherry-Ena_Trimer_, and mCherry-Ena_Tetramer_. Tubulin showed as loading control. (B) Dependence of filopodia density on GFP-actin fluorescence of cells. (C) Dependence of filopodia density on the respective mCherry-Ena construct fluorescence intensity.

**Figure 5 – figure supplement 1.**
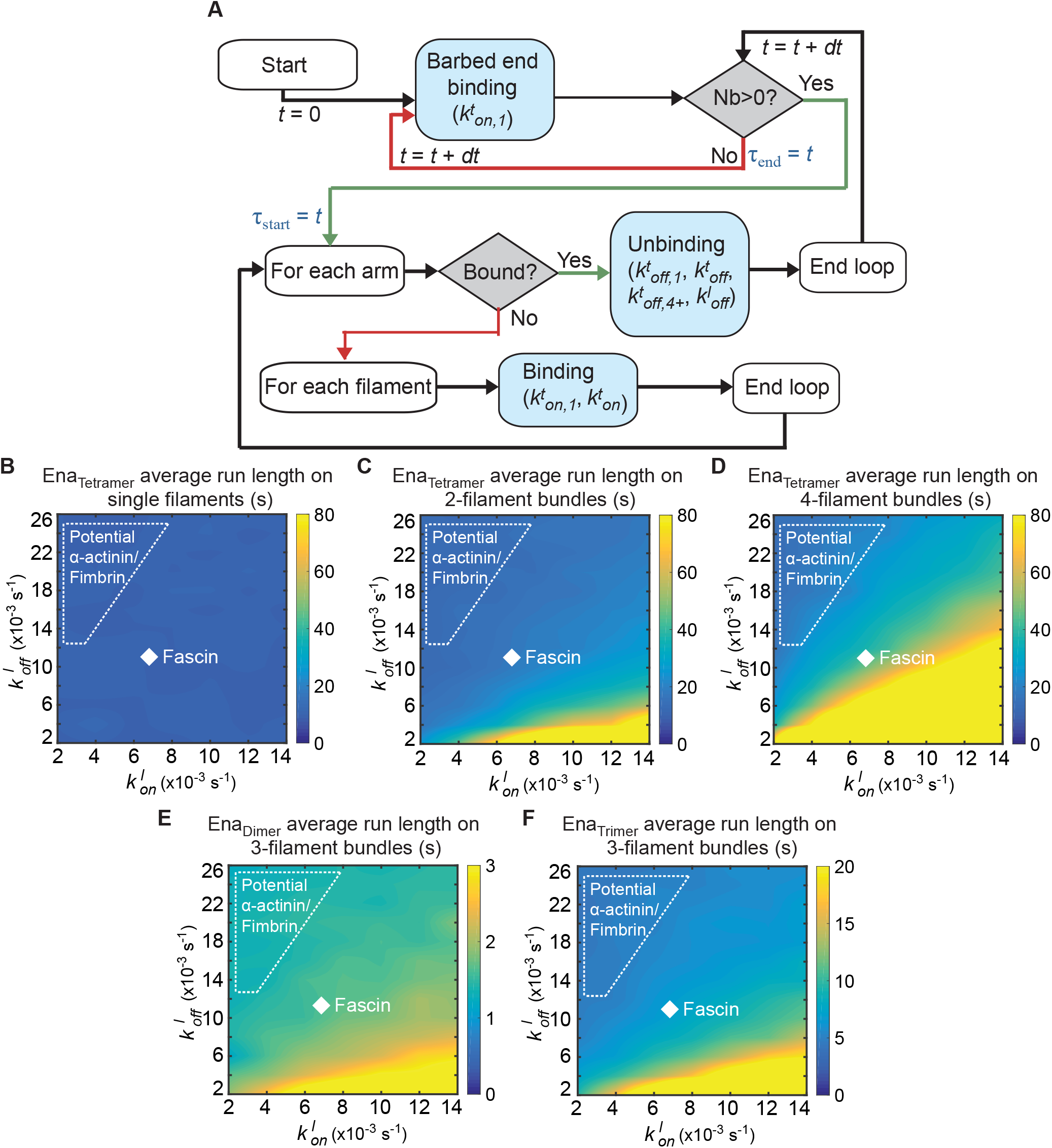
Kinetic model can explore different parameter spaces. (A) Diagram showing the key steps in the algorithm. Nb represents the number of arms bound at time *t*. τ_start_ and τ_end_ mark the start and end of Ena “molecule” binding events. The processive run length *τ* is estimated as the average of the difference *τ_end_* — *τ_start_* across several events. (B – F) Heat maps showing average Ena processive run length with systematic variations of 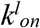 and 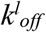 Heat maps of average processive run length for (B) single filaments (C) 2-filament bundles or (D) 4-filament bundles with four Ena arms. These are comparable to Figure 5C with 3-filament bundles. Heat maps of average processive run length for 3-filament bundles with (E) two or (F) three Ena arms.

## MOVIE LEGENDS

**Movie 1.** Ena processivity on fascin bundles (corresponds to Figure 1C-D,I,K). Spontaneous assembly of 1.5 μM Mg-ATP-actin (15% Oregon green-actin) with 15 pM SNAP(549)-EnaΔL (red) and unlabeled 130 nM human fascin visualized by two-color TIRFM. White arrowheads mark free slow-growing barbed ends, and yellow arrowheads mark fast growing barbed ends associated with Ena. Time interval between frames is 1 s.

**Movie 2.** UNC-34 processivity on fascin bundles (corresponds to Figure 2F-G). Spontaneous assembly of 1.5 μM Mg-ATP-actin (15% Oregon green-actin) with 18 pM SNAP(549)-UNC-34 (red) and unlabeled 130 nM human fascin visualized by two-color TIRFM. White arrowheads mark free slow-growing barbed ends, and yellow arrowheads mark fast growing barbed ends associated with UNC-34. Time interval between frames is 0.5 s.

**Movie 3.** Ena_Dimer_ processivity on fascin bundles (corresponds to Figure 3C,E). Spontaneous assembly of 1.5 μM Mg-ATP-actin (15% Alexa488-actin) with 50 pM MBP-SNAP(549)-EnaΔLΔCC-GCN4 (red) and unlabeled 130 nM human fascin visualized by two-color TIRFM. White arrowheads mark free slow-growing barbed ends, and yellow arrowheads mark fast growing barbed ends associated with Ena_Dimer_. Time interval between frames is 0.5 s.

## ACKNOWLEDGEMENTS

We thank Elena Solomaha of the University of Chicago BioPhysics Core Facility for performing SEC-MALS and Jonathan Winkelman for cloning VASP and UNC-34. We thank Caitlin Anderson, Katie Homa, Cristian Suarez and other members of the D.R.K. laboratory for helpful discussions. We thank the Drosophila Genomics Resource Center, supported by NIH grant 2P40OD010949, for *Drosophila* cells. This material is based upon work supported by the National Science Foundation Graduate Research Fellowship under Grant No. DGE-1144082 and DGE-1746045 (to A.J.H.), National Institute for Health’s Molecular and Cellular Biology Training Grant T32 GM007183 (to A.J.H.), National Institute for Health’s Grant RO1 GM079265 (to D.R.K.), Department of Defense Army Research Office’s MURI grant W911NF1410403 (to G.A.V. and D.R.K.), and by the University of Chicago Materials Research Science and Engineering Center, funded by National Science Foundation award DMR-1420709 (to G.A.V. and D.R.K.). Acknowledgement is made to the computational resources provided by the Research Computing Center at The University of Chicago.

